# Cellular plasticity balances the metabolic and proliferation dynamics of a regenerating liver

**DOI:** 10.1101/2020.05.29.124263

**Authors:** Ullas V. Chembazhi, Sushant Bangru, Mikel Hernaez, Auinash Kalsotra

## Abstract

The adult liver has exceptional ability to regenerate, but how it sustains normal metabolic activities during regeneration remains unclear. Here, we use partial hepatectomy (PHx) in tandem with single-cell transcriptomics to track cellular transitions and heterogeneities of ~22,000 liver cells through the initiation, progression, and termination phases of mouse liver regeneration. Our results reveal that following PHx, a subset of hepatocytes transiently reactivates an early-postnatal-like gene expression program to proliferate, while a distinct population of metabolically hyperactive cells appears to compensate for any temporary deficits in liver function. Importantly, through combined analysis of gene regulatory networks and cell-cell interaction maps, we find that regenerating hepatocytes redeploy key developmental gene regulons, which are guided by extensive ligand–receptor mediated signaling events between hepatocytes and non-parenchymal cells. Altogether, our study offers a detailed blueprint of the intercellular crosstalk and cellular reprogramming that balances the metabolic and proliferation requirements of a regenerating liver.

## INTRODUCTION

The liver is a multi-functional organ critical for carrying out numerous metabolic, biosynthetic, and detoxification functions. Owing to its detoxification roles, the liver is frequently exposed to many hepatotoxins resulting in tissue damage and cell death. Accordingly, it has evolved a unique ability to regenerate in response to a wide range of physical and toxic injuries (Diehl and Chute, 2013; Taub, 2004), and remarkably, mammalian livers can replenish up to 70% of the lost tissue mass and functionality within weeks of surgical resection (Bangru and Kalsotra, 2020; Michalopoulos and DeFrances, 1997; Michalopoulos, 2007; 2017). However, hepatic regeneration in humans is compromised after certain xenobiotic injuries, viral infections, chronic inflammation, or excessive alcohol consumption, which can lead to fibrosis and fulminant liver failure (Cordero-Espinoza and Huch, 2018; Forbes and Newsome, 2016; Louvet and Mathurin, 2015; Richardson et al., 2007; Seitz et al., 2018). It is estimated that nearly two million people die from liver disease every year, making it a prominent cause of global morbidity and mortality (Asrani et al., 2019; Marcellin and Kutala, 2018).

As most liver injuries trigger hepatocyte death, the regenerative course is primarily devoted to replenishing the lost hepatocyte population. Several cell-fate and lineage-tracing studies have determined that—under normal circumstances—the majority of new hepatocytes are derived from pre-existing hepatocytes instead of hepatic stem cells (Font-Burgada et al., 2015; Schaub et al., 2014; Yanger et al., 2014). Intriguingly, depending on the extent of the injury, surviving hepatocytes rely on hypertrophic growth, cellular proliferation, or both to restore normal liver function (Bangru and Kalsotra, 2020; Miyaoka et al., 2012). Consequently, in order to stimulate cell division and growth, the regenerating hepatocytes undergo global alterations in gene expression, which are achieved by dynamic changes in mRNA abundance, splicing and translation (Aloia et al., 2019; Bangru et al., 2018; Hyun et al., 2020; Rychtrmoc et al., 2012; Sato et al., 2017; Wang et al., 2020; 2019; Zahm et al., 2020). Although many previous studies have focused on the proliferative capacity of hepatocytes, the exact mechanics of regeneration such as how quiescent hepatocytes transition into a proliferative state, how regenerating livers sustain normal metabolic activities as the tissue recovers from injury, or what cell-cell interactions initiate and terminate the regenerative response is unknown.

Here, we leveraged a single-cell RNA sequencing (scRNA-seq) strategy to capture all resident cell types from mouse livers and dissected their cellular heterogeneities and responses to 70% partial hepatectomy (PHx) during the initiation, progression, and termination phases of regeneration (Mitchell and Willenbring, 2008; Zheng et al., 2017). Our analyses revealed that following PHx, the transcriptomes of regenerating hepatocytes are extensively reprogrammed as they bifurcate into “metabolically hyperactive” or “proliferating” states. Using in-depth trajectory inferences, we found that after PHx, a subset of residual hepatocytes reversibly activate an early-postnatal-like gene program to support cell division and growth while a distinct population of hepatocytes upregulates their adult metabolic gene program to offset regeneration-induced deficits in liver function. Our combined analysis of gene regulatory networks and cell-cell interactions uncovered that dynamic changes in the activity of key regulons within hepatocytes—orchestrated by the activation of specific non-parenchymal cells—balance the metabolic and proliferation needs of a regenerating liver. These findings support a division of labor model wherein hepatocytes acquire alternate states to enable normal metabolic activities as the liver restores its lost tissue mass. We also identify a vast array of ligand-receptor interactions among hepatocytes, endothelial, Kupffer, stellate, and T cells that coordinate the overall time course of liver regeneration. Thus, our study offers a high-resolution view of the cellular and molecular basis of liver regeneration while providing a rich resource for the identification of genes and signaling pathways that facilitate hepatic repair in response to injury.

## RESULTS

### Cell type composition, heterogeneity and metabolic dynamics of a regenerating liver

Surgical resection of the adult mouse liver by 2/3^rd^ PHx induces rapid hyperplasia and hypertrophy in the remnant tissue, such that the liver recovers its original mass and function within seven days (Boyce and Harrison, 2008; Mitchell and Willenbring, 2008) **(Figure 1A, Figure 1—figure supplement 1A)**. Hepatocytes, which constitute the bulk of liver parenchyma are among the first cells to enter cell cycle after PHx, followed by the proliferation of other stromal cells, (Fausto et al., 2006; Su et al., 2002). By labeling new DNA synthesis with 5-ethynyl-2’-deoxyuridine (EdU) and combining it with hepatocyte nuclear factor 4-alpha (*Hnf4α*) immunostaining (Bangru et al., 2018), we detected maximal hepatocyte proliferation activity between 24 and 72 hours (h) after PHx, which peaked around 36-48h **(Figure 1B, Figure 1— figure supplement 1B)**. Therefore, to sample the cellular composition and diversity as well as profile their regenerative response at a single-cell resolution, we utilized 10x genomics-based scRNA-seq platform and studied the transcriptomes of all resident cell types isolated from mouse livers at 24, 48, and 96h after PHx or sham surgery **(Figure 1C, Figure 1—figure supplement 1C)**. In parallel, we also collected cells from postnatal day 14 (P14) livers—a midpoint between the neonatal period and weaning—and performed scRNA-seq to analyze the cellular transitions and gene programs associated with normal maturation of the liver. Single cells were isolated by two-step collagenase perfusion (Bhate et al., 2015; Li et al., 2010), followed by magnetic-activated cell sorting that allows rapid and easy removal of dead cells.

**Figure 1:**
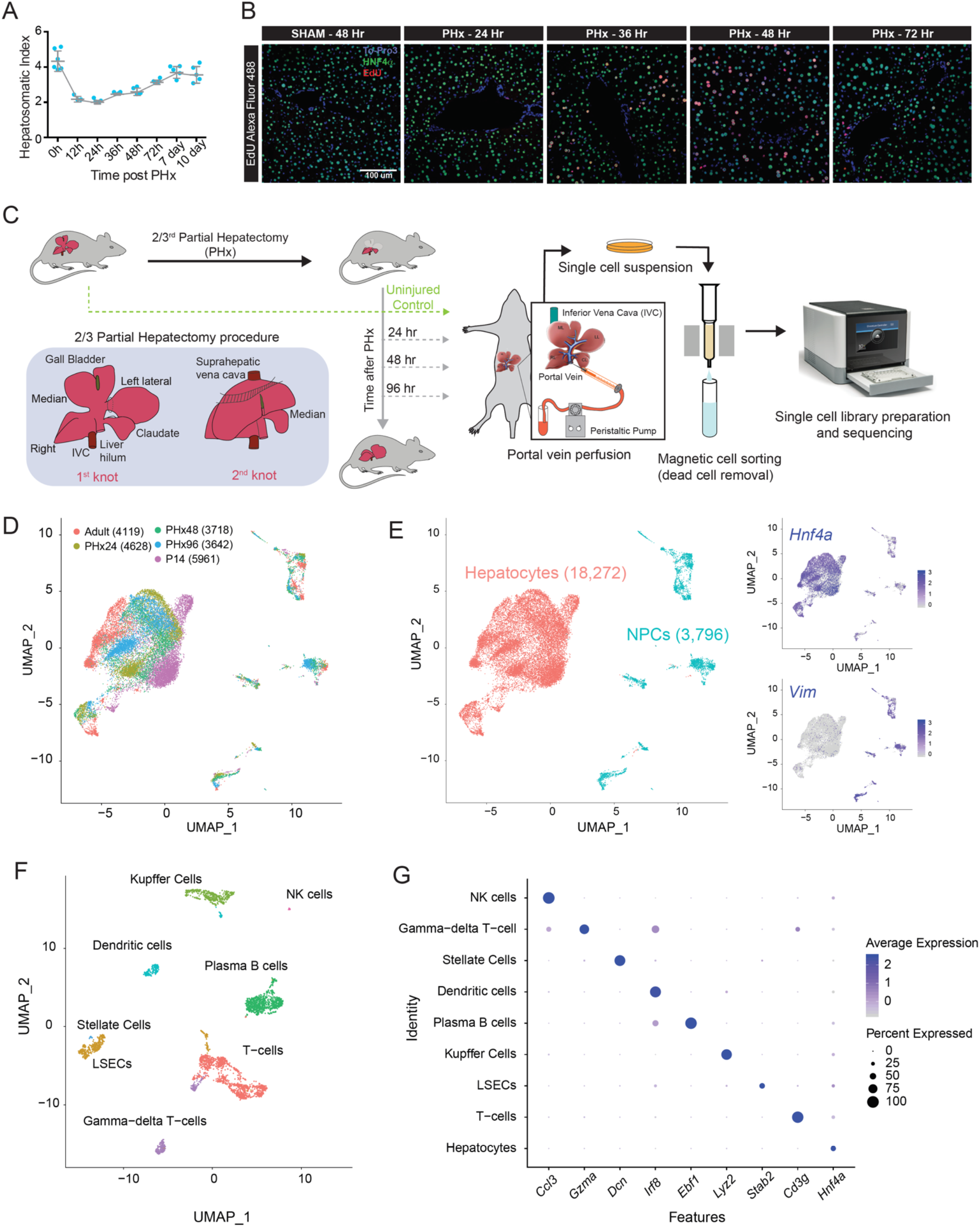
Single cell analysis of resident hepatic cell populations from immature, adult and regenerating mouse livers. **A.** Time course plot showing the restoration of liver-to-bodyweight ratio after partial hepatectomy (PHx). The liver recovers its original mass within 7 days after PHx. (n = 5 animal per group) **B.** Fluorescent imaging of hepatocyte proliferation measured by *in vivo* EdU incorporation in post-PHx and Sham livers. White arrows indicate proliferating hepatocytes (Co-labeled for HNF4a in green, incorporated EdU in red, and nuclei in blue). Images taken under 20X resolution are shown. **C.** Overview schematic demonstrating workflow for isolation of mouse liver cells for singlecell RNA sequencing (scRNA-seq). Portal vein perfusion of collagenase containing buffer was used to isolate single liver cells from P14 pups, sham adults, as well as mice at 24 h, 48 h and 96 h after 2/3^rd^ PHx. Single cell library preparation was performed with whole cell suspensions individually for each mouse using 10X Chromium Single Cell 3’ Reagent Kit (V3 chemistry) after Magnetic-activated cell sorting to remove dead cells. The inset details our PHx procedure, showing the position of two knots before excision of the respective liver lobes. **D.** Combined UMAP clustering of all 22,068 cells identified after QC cutoffs and batch correction. Cells are colored by the batch of origin and the total number of cells identified from each batch are given in brackets. **E.** Identification of hepatocyte and Non-parenchymal cell (NPC) subpopulations. Seurat based clustering followed by marker gene analysis revealed broad epithelial and non-epithelial cellular identities. Feature plots shown as insets show higher expression of expression of *Hnf4a* (a hepatocyte marker) and *Vim* (a non-epithelial marker) specially in populations identified as hepatocytes and NPCs respectively. **F.** Combined UMAP clustering analysis of all NPC subpopulation. Analysis of known cell-specific marker genes revealed the identities of each cluster as a distinct NPC cell type. **G.** Dot plot showing expression of cell-type specific marker genes for each cell-type identified in the integrated dataset. The size of each dot encodes the percentage of cells expressing that marker within the identity class (cell type) and the color encodes the average expression of a gene among all cells within that class.

After stringent quality control and normalization **(Figure 1—figure supplement 2A-C)**, we captured a total of 22,068 cells that were evenly distributed among the five time points and had a mean UMIs of 2148 and a median of 1097 genes per cell **(Figure 1D—figure supplement 2B)**. A higher fraction of reads from the mitochondrial genome is often associated with low quality or dying cells (Ilicic et al., 2016). Because hepatocytes possess a very high mitochondrial content (MacParland et al., 2018; Weibel et al., 1969), we used a relatively higher percentage of mitochondrial-read threshold (30%) in our analysis. To allow cross-sample comparisons, we integrated datasets from all samples and corrected their batch effects using the BEER algorithm (Zhang et al., 2019). Cell-type identity was assigned based on the top differentially expressed genes and from the previously identified lists of canonical cell-type-specific markers (MacParland et al., 2018; Xiong et al., 2019). Next, a graph-based clustering was performed to group cells according to their gene expression profiles. Visualization of the integrated dataset using Uniform Manifold Approximation and Projection (UMAP) classified 18,272 hepatocytes and 3,796 non-parenchymal cells (NPCs) based on their relative expression of *Hnf4*α, and the mesenchyme-derived cell marker Vimentin (*Vim*), respectively **(Figure 1D, E)**. The NPC population was further resolved into eight distinct clusters that represented liver sinusoidal endothelial cells (LSECs), stellate cells, Kupffer cells, dendritic cells, T-cells, γ-δ T-cells, Plasma B cells and NK cells **(Figure 1F, G)** and **(Figure 1—figure supplements 3A, B, and 4)**. We could not identify any cholangiocytes, and we attribute this absence to their relatively low numbers in quiescent and PHx-induced regenerating livers. Of note, previous single-cell studies that analyzed NPCs also failed to capture cholangiocytes after random sampling and had to apply additional sorting strategies to enrich for these small biliary epithelial cell (BEC) populations (Aizarani et al., 2019; Pepe-Mooney et al., 2019; Xiong et al., 2019).

Metabolic adaptation due to hepatic insufficiency is a hallmark of liver tissue renewal and regeneration (Caldez et al., 2018; Huang and Rudnick, 2014; Locasale and Cantley, 2011). Accordingly, we assessed changes in the strength of metabolic pathways among hepatocytes isolated from naïve and regenerating (PHx) livers. Several pathways were coordinately downregulated at 24 and 48h after PHx, evident from a global shift in their pathway score distribution **(Figure 2A, B)**. Whereas the gene sets belonging to fatty acid, lipid, and amino acid metabolism showed the most drastic decrease through the initiation (24h after PHx) and progression (48h after PHx) phases of regeneration, other pathways such as glycolysis, gluconeogenesis, and pentose monophosphate shunt were only subtly muted **(Figure 2A).** Notably, the dampening of liver metabolism was transient and largely reversed in the termination phase (96h after PHx), revealing the dynamic nature of such changes evoked during regeneration. Moreover, while assessing the strength of different pathways/gene sets, we found striking trends wherein at times when biosynthesis, detoxification, complement/coagulation and other secretory functions associated with mature hepatocytes were downregulated, the pathways related to cell cycle, proliferation, and growth such as ribosome biogenesis as well as RNA processing, splicing, and translation were upregulated **(Figure 2B).** Interestingly, although the cell cycle and growth-related pathways had predominantly switched off at 96h after PHx, some of the adult hepatocyte functions were not yet fully restored by this time. These results are in line with those of recent reports that explored how cell division and energy metabolism intersect in support of liver regeneration (Bangru et al., 2018; Caldez et al., 2018; Wang et al., 2018; 2019); and they highlight the metabolic flexibility of regenerating hepatocytes as they repopulate the liver to restore its lost mass and function.

**Figure 2:**
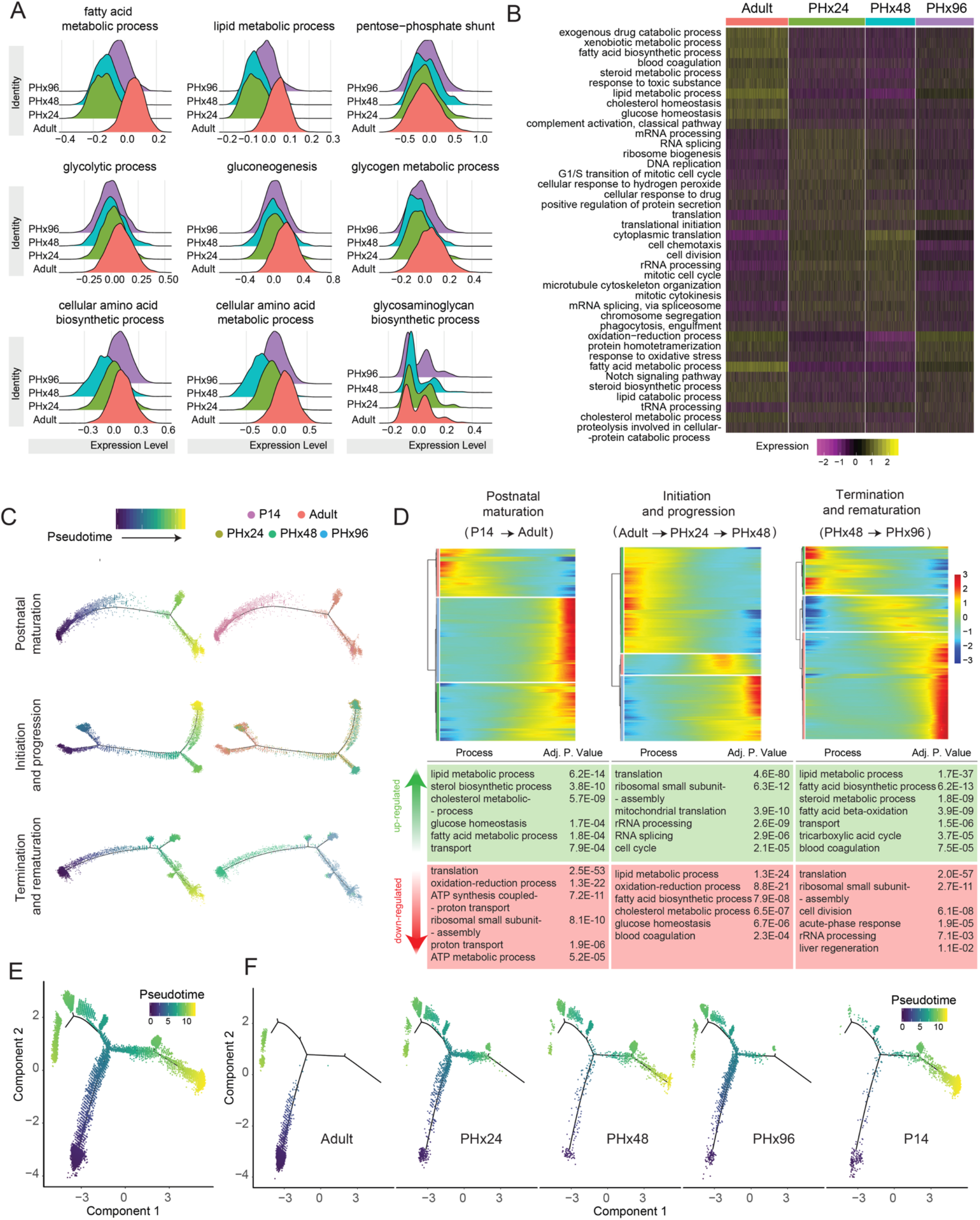
Specific hepatocyte population reversibly reprograms to an immature post-natal like state during regeneration. **A.** Ridge plots showing relative scoring on hepatocyte subpopulation using Seurat3.1 demonstrate extensive rewiring of metabolic genes during regeneration. Relative scores were computed based on the lists of genes for each pathway obtained from the Rat Genome Database (RGD). **B.** Heatmap showing relative scores of top differentially-regulated metabolic pathways. **C.** Pseudotime plots demonstrating cellular trajectories during postnatal maturation (including P14 and adult hepatocytes), initiation and progression (including adult, PHx24 and PHx48 hepatocytes), and termination and rematuration (including PHx48 and PHx96 hepatocytes). Single cell trajectories were constructed and pseudotime values calculated using Monocle 2. Trajectories are colored by pseudotime (left) and sample identity (right). **D.** Heatmaps representing modules of genes that co-vary along the pseudotime during postnatal maturation, initiation–progression, and termination–rematuration phases. DAVID based gene ontology (GO) analysis revealed reversible reprogramming of developmentally regulated gene expression programs essentially revert postnatal maturation, and this is followed by transitions that reinstate mature hepatic program. Top upregulated and downregulated GO terms are described below the respective heatmaps. **E.** Pseudotime plot indicating cellular trajectories of hepatocytes from all samples. Single cell trajectories were constructed and pseudotime values were calculated using Monocle 2. Trajectories are colored by pseudotime. **F.** Pseudotime plots showing distribution of each sample along combined cellular trajectories shown in (C). The adult and P14 hepatocytes present strikingly distinct distribution along the trajectory, however, the distribution shifts towards P14 at PHx24-48 and back towards the adult at PHx96.

### Pseudo-temporal arrangement of hepatocytes unveils cellular transitions in regeneration and reprogramming to a postnatal-like state

To map the cellular transitions as hepatocytes move from a quiescent to proliferative state and back, we performed pseudotime ordering of hepatocytes and built their individual trajectories. We used DDRTree implementation in Monocle 2, wherein top differentially expressed genes among hepatocytes collected from different time points were used for ordering cells (Qiu et al., 2017b). We constructed discrete cell-state trajectories for **1)** the normal postnatal maturation (P14➔Adult), **2)** the initiation–progression phase of regeneration (Adult➔PHx24➔PHx48), and **3)** the termination–rematuration phase of regeneration (PHx48➔PHx96) **(Figure 2C)**. For both postnatal maturation and termination–rematuration trajectories, the cells from respective time points yielded distinct branches with a few transitioning cells **(Figure 2C, top & bottom panels)**. Conversely, for the initiation–progression phase trajectory, whereas a large number of regenerating hepatocytes diverged away from the adult state, a significant portion retained their mature identity, revealing a distributive model of regeneration **(Figure 2C, middle panel)**.

Next, to identify the functional pathways changing within individual trajectories, we determined the expression dynamics of top 2000 genes that vary as a function of progress through the pseudotime. Along the postnatal maturation trajectory, the expression of genes encoding RNA processing, ribosome assembly and translational regulation factors declined first, followed by simultaneous increase in the expression of genes associated with various adult hepatocyte functions **(Figure 2D, left panel)**. Recently, bulk transcriptome analyses revealed that in response to toxin-mediated injury, regenerating hepatocytes reprogram to an early postnatal-like state (Bangru et al., 2018). But, it is unclear whether regeneration after PHx involves a similar reprogramming event. We observed that along the initiation–progression trajectory of regeneration—opposite to postnatal maturation—many metabolic genes in hepatocytes were downregulated prior to the upregulation of genes encoding cell cycle, RNA processing, and translation regulation factors. Particularly, the genes involved in ribosome biogenesis and assembly were activated only briefly along the pseudotime, underscoring that a temporary boost in protein synthesis is needed to prime hepatocytes for cell cycle re-entry **(Figure 2D, middle panel)**. Meanwhile, along the termination–rematuration trajectory—similar to postnatal maturation—downregulation of cell cycle, RNA processing, and translation related genes preceded a synchronous increase in the expression of various metabolic and biosynthetic genes **(Figure 2D, right panel)**.

It has been postulated that most hepatocytes can re-enter the cell cycle and proliferate after 2/3^rd^ PHx (Michalopoulos, 2007). To determine if all hepatocytes after PHx reprogram to a postnatal-like state and proceed simultaneously towards the proliferative trajectory, we performed pseudotime ordering of cells from all time points (i.e., P14, Adult (sham), PHx24, PHx48 & PHx96) **(Figure 2E).** We found that while the P14 and adult hepatocytes resided on separate arms of the trajectory, regenerating hepatocytes were distributed among all three arms **(Figure 2F)**. Interestingly, the majority of hepatocytes at 24h after PHx migrated away from their adult position at the beginning of pseudotime and occupied a position around the branch point **(Figure 2F)**. At 48h after PHx, a large number of hepatocytes inhabited the far right arm of the trajectory overlapping with the position of P14 hepatocytes. However, at 96h after PHx, most hepatocytes had left the P14 state returning back to their initial adult state **(Figure 2F)**. Thus, our pseudotime analysis captured the remarkable cellular plasticity exhibited by hepatocytes as they progress through different stages of regeneration. Intriguingly, at all times, a considerable population of cells remained adult-like and occupied their original position on the pseudotime axis, indicating that not all hepatocytes are reprogrammed after PHx **(Figure 2F)**. Together, these findings illustrate that reversible postnatal-like reprogramming facilitates hepatocytes in transitioning from a quiescent to proliferative state and back. The dampening of mature hepatocyte characteristics followed by a transient increase in global protein synthesis is likely a pre-requisite for cell cycle re-entry and to jump-start the regenerative process.

### Division of labor balances the metabolic and proliferation activities of regenerating hepatocytes

The DDRTree algorithm resolved our pseudotime trajectory into nine distinct hepatocyte populations (HEP1–HEP9) **(Figure 3—figure supplement 1A).** Consequently, to gain a deeper insight into the HEP1–HEP9 populations, we determined their gene expression profiles along the pseudo-temporal trajectory. We found that HEP1 and HEP2 populations were enriched for genes expressed in immature hepatocytes and de-enriched for genes expressed in mature hepatocytes **(Figure 3—figure supplement 1B)**. Contrary to HEP1 and HEP2, the HEP4 and HEP7 populations were enriched for genes expressed in mature and de-enriched for genes expressed in immature hepatocytes. HEP3, HEP5–6, and HEP8–9 populations showed intermediate enrichment for immature and de-enrichment for mature gene expression **(Figure 3—figure supplement 1B)**. Based on these transcriptome signatures, we categorized HEP1– HEP9 populations into four broader clusters. The sham adult HEP4 cluster was designated as the “quiescent state”. The cluster near the branch point emerging after PHx and comprising HEP3, HEP5–6, and HEP8–9 populations was termed as the “transition state”. Further, the cluster formed by HEP1 and HEP2 populations was marked as the “proliferative state”, whereas the cluster formed by the HEP7 population was designated as the “metabolically hyperactive state” **(Figure 3A, B)**.

**Figure 3:**
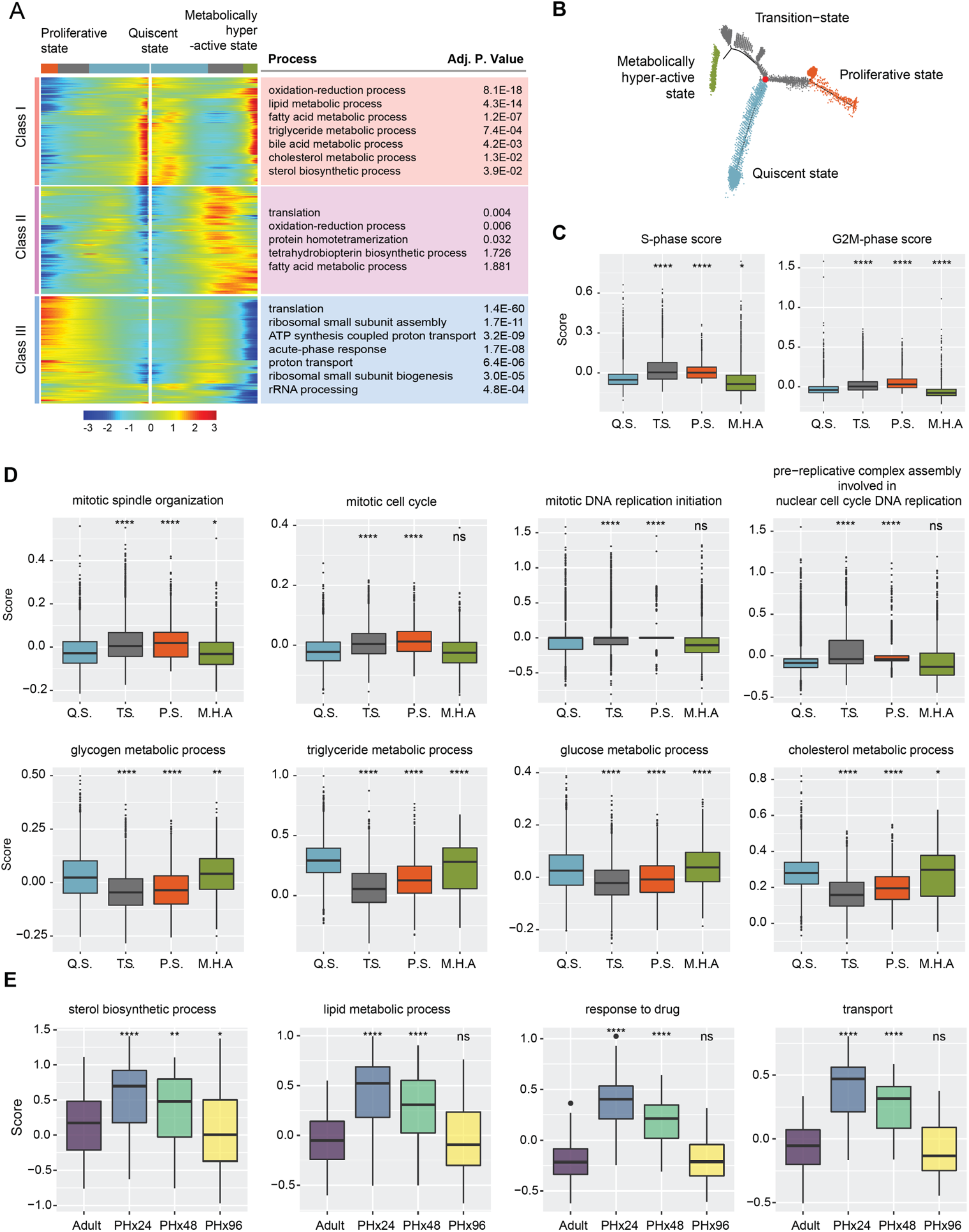
Bifurcation of hepatocyte trajectory during regeneration produces hepatocytes enriched with complementary functions in proliferation and metabolism. **A.** Heatmap showing bifurcating of gene expression programs executed along the pseudotime after branching. Top GO terms enriched in each class of genes are listed with their corresponding adjusted P-values. **B.** Trajectory demonstrating the three distinct states of hepatocytes. Branch point under evaluation is shown in red. **C.** Box plots demonstrating cell cycle phase scores calculated from Seurat3.1 for hepatocytes belonging to different states. Q.S. is quiescent state, T.S. is transition state, P.S. is proliferating state, and M.H.A. is metabolically hyperactive state. P-values were derived from a parametric t-test (unpaired). *p ≤ 0.05, ****p ≤ 0.0001 **D.** “Proliferating’ and ‘metabolically-hyperactive’ states uniquely upregulate proliferation-or metabolism-related functions, respectively. Box plots showing relative scoring of indicated pathways in hepatocytes belonging to different states. P-values were derived from a parametric t-test (unpaired). *p ≤ 0.05, **p ≤ 0.01, ****p ≤ 0.0001, ns: p > 0.05. **E.** Metabolically-hyperactive state transiently upregulates metabolism-related functions during regeneration. Violin plot showing time point based scoring of hepatocytes from the metabolically hyperactive state for the indicated pathways. P-values were derived from a parametric t-test (unpaired). *p ≤ 0.05, **p ≤ 0.01, ****p ≤ 0.0001, ns: p > 0.05.

The quick emergence of the transition state following PHx suggested that it is derived from the quiescent state, after which it bifurcates towards the proliferative or metabolically hyperactive states. To further study how gene expression diverges after PHx and generates discrete clusters of proliferative and metabolically active hepatocytes, we used BEAM module analysis within the Monocle 2 pipeline. We identified genes changing along different arms of the DDRTree, and these were grouped into three main classes **(Figure 3A)**. The class I contained genes that were highly expressed in the quiescent state, downregulated as the bifurcation point was approached, and upregulated again in the metabolically hyperactive state. In contrast, classes II and III contained genes that were poorly expressed in the quiescent state, upregulated as cells progressed through the transition point, but were then reciprocally up or downregulated in the proliferative and metabolically hyperactive states, respectively. Upon further gene set enrichment analysis, we confirmed that the metabolically hyperactive hepatocytes showed overrepresentation of functional categories related to biosynthesis and metabolism **(Figure 3A, D)**. Conversely, hepatocytes associated with the proliferative state showed an overrepresentation of cell cycle and growth related functional categories, including DNA replication, and mitosis **(Figure 3A, C, D)**. Consistent with these results, recent histological analysis of regenerating mice livers after PHx detected intertwined sets of hepatocytes that clearly segregate according to elevated glycogen content with low mitotic activity or reduced glycogen content with high mitotic activity (Minocha et al., 2017).

Having demarcated the four cellular states along the trajectory, we reasoned that if metabolically hyperactive hepatocytes do actually compensate for any temporary deficits in liver function while other hepatocytes proliferate, they should exhibit a surge in the expression of metabolic genes after PHx. To test for this possibility, we analyzed the cells of metabolically hyperactive state in relation to their time point of origin. Interestingly, relative to sham adults, metabolically active hepatocytes at 24h after PHx showed significantly higher expression of biosynthetic, metabolic, detoxification, and transport related genes, which started to decline at 48h and were essentially reversed by 96h after PHx **(Figure 3E)**. Collectively, these data support a division of labor model—wherein after PHx—a subset of residual hepatocytes acquire the metabolically hyperactive state that upregulates its adult gene program to counteract regeneration-associated deficits in liver function.

### Rewiring of gene regulatory networks activates cell state transitions during regeneration

To delineate the gene regulatory networks (GRNs) that might stimulate various cell state transitions in regeneration, we used single-cell regulatory network inference and clustering (SCENIC) pipeline (Aibar et al., 2017) on our scRNA-seq data (see methods). SCENIC computes the activity of transcription factors from individual cells by integrating co-expression data with transcription factor motif enrichment analysis, generating a “regulon”, which refers to an expressed transcription factor and all of its co-expressed target genes. We obtained the regulon activities using AUCell, which ranks targets of each regulon among the expressed genes in each cell, yielding a regulon-by-cell activity matrix. The overarching function of SCENIC is to create regulon-driven clusters that are generated from the regulon-activity matrix through binarizing (by thresholding) the original AUCell score. We, however, discerned that instead of binarizing, maintaining the full AUCell score improves the inferences performed on the data.

We hypothesized that the reimplementation of certain developmental GRNs might drive cellular transitions during regeneration. To test this hypothesis, we analyzed the AUCell score activity matrix of individual hepatocytes acquired from all time points (i.e., P14, Adult, PHx24, PHx48 & PHx96). Upon visualization of cell clustering from UMAP—built from the AUCell scores—we noted that hepatocytes from specific time points grouped together, underscoring a high degree of similarity in their regulon activity (**Figure 4A, B**). Importantly, the P14 and adult stage hepatocytes formed distinct non-overlapping clusters at far ends of the UMAP plot, representing clear differences in their GRNs. The hepatocytes from 24h and 48h after PHx, however, clustered adjacent to P14 and away from the adult stage, demonstrating that similar regulons are active during postnatal development and initiation–progression phases of regeneration. Interestingly, very few PHx96 hepatocytes overlapped with P14, PHx24, or PHx48 time points (**Figure 4A, B**). Instead, they were clustered around adult cells, which indicates that in the termination phase, hepatocytes restore their mature adult-like regulon activity. Based on these findings, we conclude that in order to reprogram gene expression after PHx, the liver cells, in part, redeploy the same GRNs that are utilized for physiologic growth during the postnatal period of liver development.

**Figure 4:**
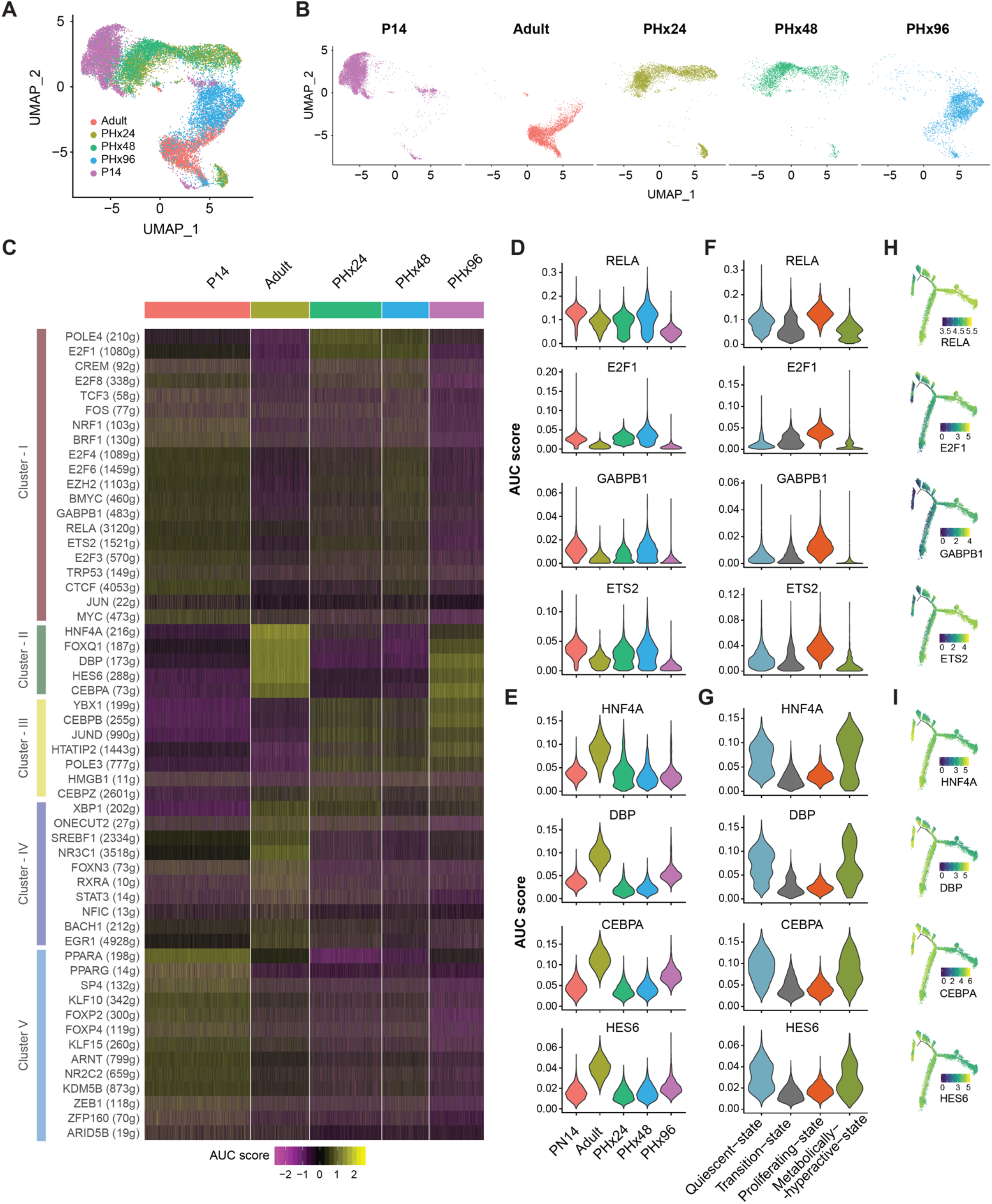
Gene regulatory networks are rewired to a postnatal-like-state during regeneration. **A.** UMAP clustering of all hepatocytes based on the AUC scores for each regulon calculated with SCENIC. Cells are colored according to the sample of origin. **B.** AUC score based UMAP clustering, grouped according to the sample of origin. Cells are colored according to sample of origin. Adult and PHx96 hepatocytes cluster together, whereas PHx24 and PHx48 hepatocytes cluster together with P14 hepatocytes. **C.** Heatmap depicting the activities of different regulons that show time point-dependent variations. **D.** Violin plot showing distribution of AUC scores for RELA, E2F1, GABP1 and ETS2 regulons across hepatocytes from each time point demonstrating their high activity in PHx24, PHx48 and P14 hepatocytes. **E.** Violin plots showing distribution of AUC scores for HNF4A, DBP, CBPA and HES6 regulons across hepatocytes from each time point demonstrating their high activity in adult and PHx96 hepatocytes. **F.** Violin plots showing distribution of AUC scores of representative regulons across hepatocytes showing their upregulation in proliferative state. **G.** Violin plots showing distribution of AUC scores of representative regulons across hepatocytes showing their upregulation in quiescent and metabolically active states. **H.** Pseudotime plots of hepatocyte cellular trajectories colored by the AUC scores of representative regulons showing high activity in the proliferative state. **I.** Pseudotime plots of hepatocyte cellular trajectories colored by the AUC scores of representative regulons showing high activity in quiescent and metabolically active states.

Next, we explored regulon activities distinguishing the cellular features of normal and regenerating hepatocytes. Our goal was to identify GRNs that are selectively activated during the initiation–progression and the termination–rematuration stages of regeneration. We reasoned that regulon activities of the initiation–progression GRNs would typically be low in adult hepatocytes, stimulated at PHx24 and PHx48, and muted again at PHx96. Correspondingly, regulon activities of the termination–rematuration GRNs would normally be high in adult hepatocytes, muted at PHx24 and PHx48, and stimulated again at PHx96. Indeed, within the 56 differentially active regulons, we detected many that fit these criteria **(Figure 4C, Figure 4— figure supplement 1A-D)**. For instance, RELA, E2F1, GABPB1, ETS2 regulons were active in hepatocytes from P14 and the initiation–progression stage of regeneration, but were turned off in adult hepatocytes and at the termination–rematuration stages **(Figure 4C cluster I, 4D)**, This indicates that these regulons likely play dual roles in regulating the hepatocyte hyperplastic response — i.e., in normal liver development and following an injury in adult animals. In line with our results, previous studies have found that a rapid increase in the expression and/or DNA binding activity of NF-kB (RELA and RELB), E2Fs (E2F1, 3, 4, 6 and 8), AP-1 (JUN and FOS), POLE4, TRP53, MYC, CREM, and ETS (ETS2, GABPB1) family of transcription factors is involved in the initiation of stress signaling, oxidative stress, DNA replication/repair, and cell-cycle entry at the early stages of liver regeneration (Baena et al., 2005; Beyer et al., 2008; Bhat et al., 1987; Chaisson et al., 2002; Colak et al., 2020; Iimuro et al., 1998; Inoue et al., 2002a; Kelley-Loughnane et al., 2002; Kurinna et al., 2013; Servillo et al., 1998; Sladky et al., 2020; Stepniak et al., 2006; Su et al., 2002; Westwick et al., 1995; Wu et al., 2013; Yang et al., 1991; Zellmer et al., 2010).

In contrast, HNF4A, DBP, CEBPA, and HES6 regulons were highly active in adult hepatocytes, muted at P14 and the initiation–progression stages, but then reactivated at the termination–rematuration stage **(Figure 4C cluster II, 4E)**, pointing towards their role in the termination of liver regeneration. The function of hepatocyte nuclear factor 4A (HNF4A), a nuclear receptor, in hepatocyte differentiation is well established (Kyrmizi et al., 2006; Parviz et al., 2003) — as it directs the expression of gene programs involved in xenobiotic, carbohydrate, and fatty acid metabolism as well as in bile acid synthesis, blood coagulation, and ureagenesis (Hayhurst et al., 2001; Inoue et al., 2002b; 2006a; 2004; 2006b; Nishikawa et al., 2015; Schrem et al., 2002). Previous studies have described HNF4A’s anti-proliferative effects in hepatocytes (Bonzo et al., 2012; Hatziapostolou et al., 2011; Walesky et al., 2013), and more recently, it was found that HNF4A is indispensable for terminating regeneration after PHx (Huck et al., 2019). Consistent with our regulon activity scores, HNF4A protein levels diminish rapidly after PHx, and this initial decrease followed by re-expression is needed for hepatocytes to timely enter and exit the cell cycle and to re-establish mature liver functions once regeneration is complete (Huck et al., 2019).

Like HNF4A, dynamic and temporally regulated activities of the CAAT/Enhancer-Binding Proteins (CEBPs) are critically important for coordinating gene expression changes through the shifting phases of regeneration (Greenbaum et al., 1995; 1998; Jin et al., 2015). CEBPA and CEBPB are basic region leucine zipper containing transcription factors that act as homo or heterodimers and bind similar DNA sequences (Osada et al., 1996). Interestingly, we found that CEBPA and CEBPB regulon activities exhibit an opposing pattern through the initiation– progression and termination–rematuration stages of regeneration **(Figure 4C, E, Figure 4— figure supplement 1B)**. CEBPA regulon activity—comprising metabolic and liver homeostatic genes—is high in pre-PHx adult hepatocytes, suppressed at PHx24 and PHx48, then enhanced in PHx96 hepatocytes. Conversely, CEBPB regulon activity—comprising many acute phase and cell cycle related genes—is low in pre-PHx adult hepatocytes, but rapidly up-regulated after PHx. Of note, CEBPA and CEBPB usually bind the same genomic locations in hepatocytes (Jakobsen et al., 2013), except with divergent temporal patterns during regeneration (Kuttippurathu et al., 2017). These dynamic shifts in genomic occupancies are tightly linked to the changes in the relative ratio of CEBP proteins such that a high CEBPA:CEBPB ratio promotes binding to *cis*-regulatory sequences boosting metabolic and suppressing acute phase response genes, whereas a low ratio directs binding to sequences that repress metabolic and activate cell cycle and acute phase related genes (Jakobsen et al., 2013). Although direct roles of D-box binding protein (DBP, a circadian PAR bZIP transcription factor) or HES family of basic helix-loop-helix transcription factor 6 (HES6) in liver regeneration are not yet explored, their regulon activity patterns **(Figure 4E)** hint both as likely termination factors.

We also recognized several regeneration-specific regulons that became activated after PHx but were otherwise inactive in postnatal development **(Figure 4C cluster III, Figure 4— figure supplement 1A)**. Intriguingly, some developmental regulons that were redeployed following PHx maintained their activated/deactivated patterns up to 96h, indicative of their extended role in regeneration **(Figure 4C cluster IV, Figure 4—figure supplement 1B)**. Finally, we also detected some development-specific regulons that were not redeployed either in the initiation–progression or in the termination–rematuration stages of regeneration **(Figure 4C cluster V, Figure 4—figure supplement 1C)**, suggesting that a portion of the genetic machinery critical for physiological liver growth and development is dispensable for regeneration. Hence, a variety of GRNs positively and negatively impact hepatocyte proliferation, and dynamic utilization of transcription factors instructs these regulons to synchronize the timely initiation, progression, and termination of liver regeneration.

We next studied the correlation of regulon activities with the pseudo-temporal transition of hepatocytes, and the four cellular states described earlier **(Figure 4F-I, Figure 4—figure supplement 2)**. By overlaying AUCell scores along the pseudotime trajectory, we noticed that RELA, E2F1, GABPB1, and ETS2 regulons—which were active in hepatocytes from P14 and initiation–progression stages of regeneration—displayed significantly higher activity in the proliferative state relative to quiescent, transition, or metabolically hyperactive states **(Figure 4F, H)**. In contrast, HNF4A, DBP, CEBPA, and HES6 regulons—all of which promote mature functions—were much more active in quiescent and metabolically hyperactive states relative to transition or proliferative states **(Figure 4G, I)**. Collectively, these data indicate that underlying changes in regulon activities are critical for determining hepatocyte identities and cell state transitions, which help preserve the highly specialized liver functions while the regenerating tissue balances its metabolic and proliferation needs.

### Transitions in hepatocyte states dictate the dynamics of intercellular signaling during regeneration

Next, we explored the dynamics of potential cell-cell communication networks at different stages of regeneration. We first inspected the cell type-specific RNA expression of various ligands in the liver secretome and their corresponding receptors, as previously described (Xiong et al., 2019). Our analysis mapped numerous unique clusters of ligand-receptor pairs with cell-type-specific expression patterns, which highlights the distinct roles of hepatocytes and NPCs in shaping intrahepatic signaling topologies (**Figure 5A)**. Among NPCs, we noticed that LSECs and stellate cells comprised the largest clusters, underscoring their predominant roles in cell signaling **(Figure 5A, left)**. Significant differences in ligand-receptor expression profiles were also detected among hepatocytes belonging to different states, which indicates the remodeling of interaction landscapes with cell state transitions **(Figure 5A, right)**.

**Figure 5:**
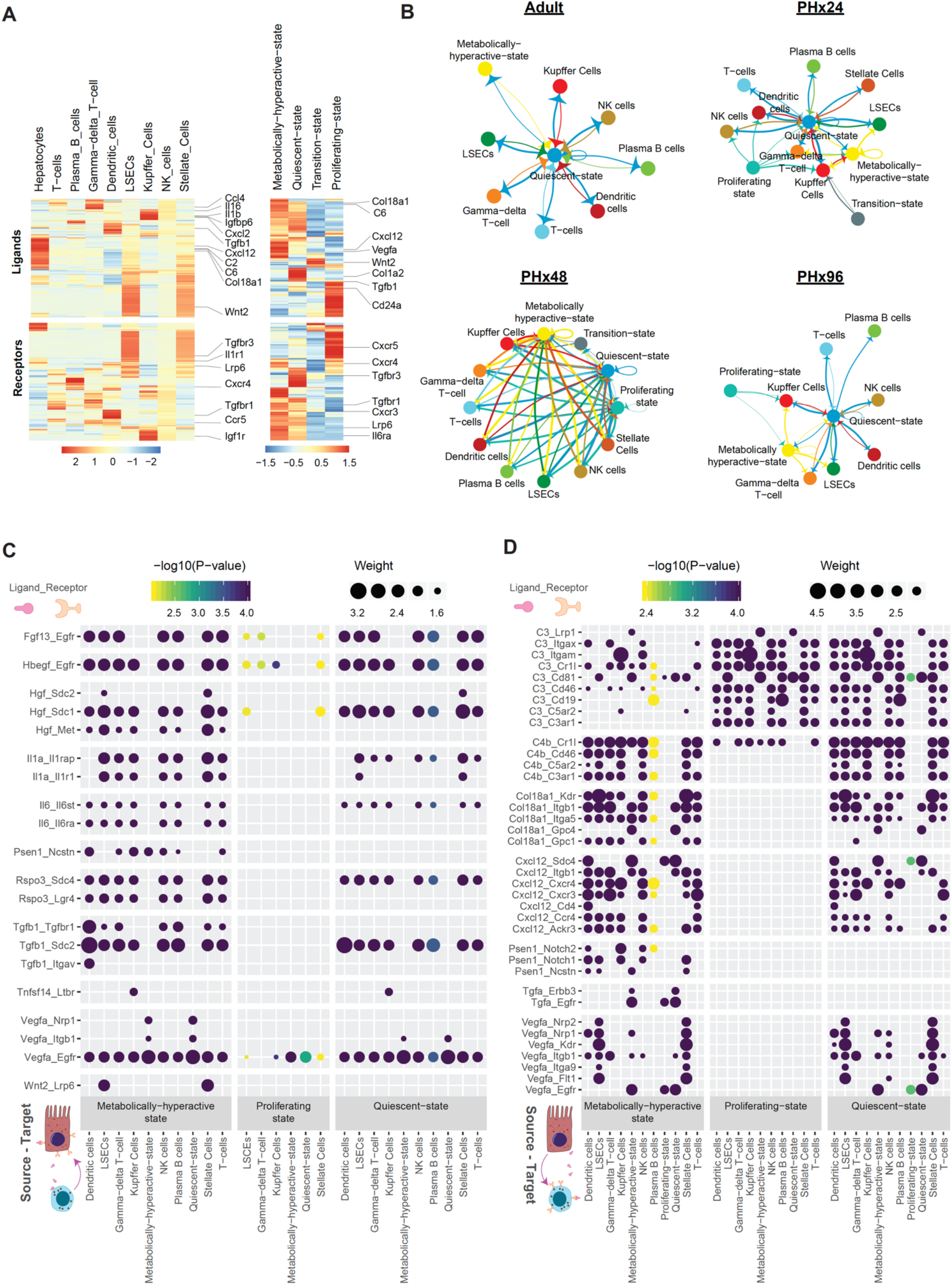
Cell-cell interactions landscape dynamics during post-PHx regenerative response. **A.** Heatmap showing expression of various ligand molecules and cellular receptors from different liver cell types (left) and from hepatocytes belonging to different cell states (right). **B.** Network diagrams showing cell-cell interactions indicated by arrows (edges) pointing in the source to target direction. Thickness indicates the sum of weighted paths between populations, and the color of arrows corresponds to the source. Network diagrams for Adult, PHx24, PHx48 and PHx96 are shown. **C.** Dot plot of representative inbound signals to hepatocytes at PHx48. Size of each dot indicates the weight of the corresponding ligand-receptor interaction and the color indicates negative log_10_ P-value of the source-to-target interaction. Empirical p-values were calculated and Benjamini-Hochberg correction was performed. **D.** Dot plot of representative outbound signals from hepatocytes to various liver cells at PHx48.

Particularly, we noted that many signaling molecules with established roles in liver regeneration—such as cytokines, chemotactic factors, secreted matrix proteins, growth factors, adhesion molecules, and mitogens—originate from specific cell types (**Figure 5—figure supplement 1A-C)**. Expression of *Wnt2*, for instance, was predominantly seen in LSECs (**Figure 5—figure supplement 1C)**, reaffirming results from previous studies of liver regeneration following PHx and acute CCl_4_ toxicity (Ding et al., 2010; Preziosi et al., 2018; Zhao et al., 2019). Importantly, pseudotime ordering revealed that upregulation of *Wnt2* and *Hgf* expression correlates with the transition of LSECs to an activated state (**Figure 5—figure supplement 2A, B and C)**, and as reported earlier, associates with *VEGFR2-Id1* activity (Ding et al., 2010). Furthermore, *Wnt2* was expressed at much lower levels in Kupffer cells relative to LSECs (**Figure 5—figure supplement 1C)**, consistent with their minor role in Wnt–b-catenin signaling (Preziosi et al., 2018; Russell and Monga, 2018; Yang et al., 2014).

To study the intracellular crosstalk among hepatic cell types and how it is modified during regeneration, we constructed cell-cell communication networks (Farbehi et al., 2019) for each time point from our dataset **(Figure 5B, Figure 5—figure supplement 3)**. The edges of the network are directed from source to target cells, which express specific ligands and their corresponding receptors, respectively. The thickness of edges corresponds to weights representing fold-changes in the expression of ligands-receptor pairs (see methods). Together, this generated a weighted and directed network of potential cell-cell interactions within normal and regenerating livers. Strikingly, the cell-cell communication networks underwent significant rewiring, evoking a transient increase in overall cellular crosstalk during regeneration. We noticed a remarkable increase in interactions between hepatocytes and NPCs at PHx24 and particularly at PHx48, followed by a re-establishment of an adult-like communication network at PHx96. Hepatocytes displayed discrete profiles of interactions with other cell-types in a statedependent manner, as expected from the differences in ligand-receptor expression observed in **Figure 5A**. Throughout regeneration, quiescent hepatocytes consistently made strong inbound and outbound connections with most other cell types, whereas transition state hepatocytes were refractory to any crosstalk. The metabolically hyperactive hepatocytes exhibited an adaptable pattern, with prominent interactions at PHx48, but minimal interactions otherwise. Interestingly, the proliferating hepatocytes presented a unique interaction landscape with strong outbound connections and few to no inbound connections. It is noteworthy that cells transitioning to the proliferating state are more amenable to regenerative cues such that continued stimulation by pro-proliferative ligands would lead to excessive/uncontrolled proliferation. We postulate that the downregulation of receptors related to pro-proliferative signals might be crucial for limiting the endless proliferation of hepatocytes and facilitating their timely exit from the cell cycle.

Next, we studied the individual ligand-receptor interactions among various cell types. We constructed dot-plots for each time point demonstrating all ligand-receptor interactions with a minimum path weight of 1.5, for all significant cell-cell relationships (P*adj* <0.01) **(Supplementary data files 1-5)**. This provided comprehensive visualization of potential cellular interactions, divulging time-point specific differences in outbound and inbound signaling from/to hepatocytes. Even at PHx48, when maximal intercellular crosstalk was observed, proliferative state hepatocytes appeared to receive a distinctly low number of incoming signals **(Figure 5C, Supplementary data files 1-5)**, which matched with the lower inbound edges to these cells seen in **Figure 5B**. Contrary to this, we detected significant inbound signaling towards quiescent and metabolically-hyperactive state hepatocytes, which was mediated by several growth factors, interleukins, and the Wnt signaling pathway **(Figure 5C, Supplementary data files 1-5)**. For instance, consistent with earlier reports, our analysis predicted hepatocytes to receive prominent HGF/MET signaling from LSECs and stellate cells (Furge et al., 2000; LeCouter et al., 2003; Maher, 1993; Schirmacher et al., 1992). We did not capture an EGF-signaling network among different cell types, which is in agreement with low EGF expression in hepatic cells (**Figure 5— figure supplement 1C)** and its predominantly exogenous origin (Olsen et al., 1985; St Hilaire and Jones, 1982; St Hilaire et al., 1983). However, we detected prominent heparin binding (HB)-EGF signaling from Kupffer cells and LSECs and other NPC populations (Kiso et al., 1995; 2003). Notably, although TGF-β protein levels in hepatocytes are debatable (Bissell et al., 1995; Braun et al., 1988; Carr et al., 1989; Nakatsukasa et al., 1990), we found that *Tgfb1* RNA is abundant in regenerating hepatocytes as well as most NPCs, but without any significant autocrine TGF-β activity within hepatocytes (**Figure 5—figure supplement 1C, Figure 5C)**. As expected, we detected many known mitogenic signals inbound to hepatocytes such as *Fgf*, *Tnf* and *IL-6*, which were high between 24-48h after PHx, but had declined by PHx96 **(Supplementary data files 1-5)**.

Outbound signals from hepatocytes at PHx48 involved pathways such as *Tgfa*, *Vegfa*, collagen, complement, and chemokine signaling **(Figure 5D, Supplementary data files 1-5)**. We noticed contrasting ligand-receptor nodes corresponding to outbound signals in metabolically hyperactive and proliferating cells, indicating opposing expression of ligands between these cell types. Certain ligands like *Tgfa* produced by hepatocytes seemed to target hepatocytes themselves. This observation is supported by the previously proposed autocrine mode of mitogenic TGFa action (Mead and Fausto, 1989; Reddy et al., 1996; Webber et al., 1993). On the other hand, *Vegfa* ligands were directed more towards LSECs and stellate cells, in line with their known roles in the activation of these cell populations (Ankoma-Sey et al., 1998; Ding et al., 2010; LeCouter et al., 2003; Liu et al., 2009); whereas complement system ligands appeared to target diverse intrahepatic cell populations (DeAngelis et al., 2012; Strey et al., 2003; Thorgersen et al., 2019). The cellular interactome analysis also predicted signaling events with uncharacterized roles in liver regeneration. For instance, *Col18a1_Kdr/Itgb1/Itga5/Gpc1/4*, *Tgfa_Egfr/Errb3*, and/or *Psen1_Notch1/2* signaling events are excellent candidates to evaluate in the context of their function in emergence/stabilization of the metabolically hyperactive state of hepatocytes. Altogether, our single-cell connectomics analysis identified a vast array of ligandreceptor interactions among hepatocytes and NPCs, provided a network-level portrait of intercellular crosstalk within normal and regenerating livers, and offered the first glimpse into how cell state transitions shape the intrahepatic signaling at different stages of liver regeneration.

## DISCUSSION

Lately, single-cell transcriptomic methods are being employed to probe cellular heterogeneities of tissues and reconstruct developmental trajectories of individual organs (Haghverdi et al., 2016; Saelens et al., 2019; Schiebinger et al., 2019). In the case of the liver, this has led to the identification of previously unknown subpopulations of hepatocytes, cholangiocytes, endothelial cells, scar-associated macrophages, stellate cells, and myofibroblasts from healthy and diseased conditions (Aizarani et al., 2019; Dobie et al., 2019; Hyun et al., 2019; MacParland et al., 2018; Pepe-Mooney et al., 2019; Ramachandran et al., 2019; Xiong et al., 2019; Achanta et al., 2019; Cook et al., 2018). The studies revealed that more than 50% of hepatocyte genes follow a discrete zonated expression pattern in a liver lobule, and surprisingly, similar metabolic zonation exists for LSECs and stellate cells (Dobie et al., 2019; Halpern et al., 2018; 2017). Furthermore, the unbiased capture of different cell types from whole tissues has led to the discovery of genome-wide cell-cell communication networks (Farbehi et al., 2019; Raredon et al., 2019; Vento-Tormo et al., 2018; Xiong et al., 2019). Here, we demonstrate that scRNA-seq is also a powerful strategy to study gene expression dynamics and communication among diverse cell populations within a regenerating tissue.

By comparing transcriptome landscapes of distinct liver cell types through the physiologic and regenerative growth periods, we found that following PHx, residual hepatocytes reversibly activate an early-postnatal-like gene program to transition from a quiescent to proliferative state and back. Our analysis revealed that transient dampening of mature gene expression programs followed by a brief surge in ribosome biogenesis precedes cell-cycle activation and thus are likely required for injury-induced proliferation of hepatocytes. We further demonstrated that rewiring of developmental GRNs orchestrates cell-cycle entry during initiation of regeneration while facilitating re-maturation of the newly generated hepatocytes so they can resume their functions once regeneration is complete. Thus, controlled reactivation of the developmental and cell cycle gene programs in adult hepatocytes could serve as a potential therapeutic approach to replenish dying hepatocytes in diseased livers.

Regeneration requires simultaneous proliferation and maintenance of highly specialized cellular functions, and fittingly a regenerating liver continues to perform its crucial metabolic, biosynthetic, and detoxification roles (Bangru and Kalsotra, 2020; Michalopoulos and DeFrances, 1997; Michalopoulos, 2007; 2017). But, how the regenerating tissue sustains these specialized functions when large numbers of cells are proliferating is still a mystery. Based on our findings, we propose a division of labor model—wherein after PHx—surviving hepatocytes undergo defined cellular transitions to allow normal metabolic activities as the regenerating liver restores its original mass. Four principal observations support this model. First, our trajectory analyses captured the extraordinary cellular plasticity of hepatocytes identifying four distinct subpopulations representing the quiescent, transition, proliferative, and metabolically hyperactive states. Second, we discovered that after PHx, quiescent hepatocytes promptly adopt an intermediate transition state from where they branch into either proliferative or metabolically hyperactive states. Third, we noticed visibly divergent regulon activities of proliferative and metabolically hyperactive hepatocytes. Cells transitioning into a proliferative state silenced regulons coordinating mature hepatocyte functions while activating regulons that support cell growth and proliferation. Conversely, cells transitioning into a metabolically hyperactive state activated multiple regulons involved in biosynthetic, metabolic, detoxification, and transport-related functions. Fourth, we observed that the metabolically hyperactive hepatocytes developed transient but strong inbound and outbound connections with non-parenchymal cell types, whereas proliferating hepatocytes selectively downregulated the receptors for inbound signals. Elimination of receptors for inbound pro-proliferative signals might be important for limiting the endless proliferation of hepatocytes and enabling their timely cell-cycle exit. Altogether, these observations illustrate that dynamic shifts in regulon activities and cell-cell interactions broaden the hepatocellular plasticity to balance the metabolic and proliferation needs of a regenerating liver.

Previous studies have indicated that distinctly located pools of mature hepatocytes with progenitor-like features serve specialized roles in liver regeneration (Miyajima et al., 2014). Although hepatocytes expressing stem/progenitor-like markers such as LGR5^+^, SOX9^+^, AXIN2^+^, TERT^+^, or MFSD2A^+^ are detectable, their overall requirement for normal maintenance and renewal after acute or chronic liver damage is debatable (Font-Burgada et al., 2015; Huch et al., 2013; Lin et al., 2018; Lu et al., 2015; Pu et al., 2016; Wang et al., 2015). For instance, a series of recent reports questioned the notion of a dedicated regenerative cell population and they demonstrated that randomly distributed hepatocytes throughout the lobule repopulate the liver under both homeostatic and/or injury conditions (Chen et al., 2020; Matsumoto et al., 2020; Sun et al., 2020). Our single-cell transcriptomic data are consistent with these recent reports as we did not detect enrichment for any of these markers in the proliferating pool of hepatocytes. Instead, we found that after PHx, a subset of remaining hepatocytes dedifferentiates to an early postnatal-like state before proceeding towards the proliferative trajectory. The conflicting observations of earlier studies could have arisen due to different lineage-tracing models, methods of hepatocellular injury, and/or severity of the disease. As certainly, when the liver is severely damaged and hepatocyte proliferation compromised, other cells with progenitor-like characteristics can transdifferentiate into hepatocytes to reconstitute the liver (Lu et al., 2015; Michalopoulos and Khan, 2015; Raven et al., 2017).

Our understanding of molecular events that induce mature, quiescent hepatocytes to dedifferentiate and transition into a proliferative state is incomplete. In this study, we combined systematic analyses of gene regulatory networks and intercellular interactions via ligandreceptor signaling on a compensatory model of regeneration (PHx) in an otherwise healthy liver. In the future, it will be important to determine whether the hepatocyte subpopulations identified here reprogram similarly or differently in response to other types of periportal and/or pericentral liver injuries. This information is crucial to fully discern the regenerative mechanisms or lack thereof in the context of human liver disease. The use of cutting-edge spatial transcriptomics methods that correlate gene expression profiles with histology data from the same/consecutive tissue sections should provide additional information on the local heterogeneity of liver cell subpopulations and their roles in regeneration. Lately, single-cell studies have surveyed the changes in cell-cell communication between healthy, NASH, and fibrotic livers (Dobie et al., 2019; Xiong et al., 2019). Comparing these disease datasets to the compendium of intercellular interactions documented here would broaden our understanding of normal versus aberrant cell-cell communication networks, revealing defects in intrahepatic cell signaling that compromise regeneration in diseased livers. Such lines of investigations will not only map the signaling events that regulate hepatocellular plasticity but also help identify targets that may be leveraged to optimize hepatic repair and function after acute liver failure or in end-stage liver disease.

## MATERIALS AND METHODS

### Animal care and surgeries

Eight-twelve week-old male mice were used for all experimental procedures. National Institutes of Health (NIH) and UIUC institutional guidelines for the use and care of laboratory animals were followed, and all experimental protocols were performed as approved by the Institutional Animal Care and Use Committee at the University of Illinois, Urbana-Champaign (UIUC). Mice were housed in 12 h light-dark cycle with standard chow diet (Teklad Global 18% Protein Rodent Diet) and water provided *ad libitum*. We performed a 2/3^rd^ partial-hepatectomy (PHx) procedure adapted from previously reported protocols (Boyce and Harrison, 2008; Mitchell and Willenbring, 2008). Briefly, a bilateral subcostal abdominal incision was made on the skin of the anesthetized (continuous isoflurane inhalation, 2%) animal to expose the abdominal musculature. 4/0-silk ligatures were then placed across the superior gastric vessels positioned vertically on either side of the xiphoid process to minimize blood loss from the peritoneal wall. Following this, a bilateral incision was made on the abdominal wall to expose the liver. The left lateral and median lobes were ligated and excised. Following this, the peritoneum was closed with a continuous 5/0 silk suture, and the skin was closed using 7 mm reflex clips. The anesthesia was then removed, and the mouse allowed to recover on a pre-warmed heating pad. To minimize post-surgical discomfort, Carprofen (5 mg/kg) was administered subcutaneously as an analgesic immediately after surgery.

### Immunofluorescence staining

EdU labeling and immunofluorescence staining were carried out as described before (Bangru et al., 2018). 5-Ethynyl-2’-deoxyuridine (EdU) was administered intraperitoneally (100μg/g body weight) four hours before tissue collection to label nascent DNA synthesis. Harvested liver tissue was ixed in 10% neutral buffered formalin for 24 hours and embedded in paraffin. 5 μm thick sections were cut, deparaffinized in xylene, rehydrated and antigen retrieved in citrate buffer (10 mM sodium citrate, 0.05% Tween 20, pH 6.0). The sections were washed with wash buffer (Tris-buffered saline with 0.05% Triton X-100) and blocked for two hours at room temperature in blocking buffer (10% normal goat serum + 1% BSA in TBS). To visualize EdU-labelled DNA, Alexa Fluor 488 was conjugated to EdU using Click-iT EdU Alexa Fluor Kit (Thermo Fisher) as per the manufacturer’s instructions. Anti-HNF4A antibody (Abcam, C11F12) was then applied to the sections at 1:500 dilution and incubated overnight at 4 °C. Following this, the sections were washed with washing buffer and incubated at room temperature for one hour with Alexa Fluor 588 conjugated secondary antibody. Nuclei were then stained with ToPro3 (1 uM in PBS) for 15 mins at room temperature. CC/Mount aqueous mounting media (Sigma) was used to mount and coverslip the sections before imaging on a Zeiss LSM 710 microscope.

### Liver function tests

Retro-orbital punctures were used to collect whole blood from mice into Capiject gel/clot activator tubes. The serum was isolated by centrifugation at 8500g for 10 min and stored at −80 °C until further analysis. Serum chemistry analysis for ALT and AST levels were performed using commercial assay kits (Infinity Kits, Thermo Scientific) according to the manufacturer’s protocols. GraphPad Prism 6 was used to perform statistical analysis (One-way ANOVA) and prepare graphical representations.

### Tissue dissociation and isolation of liver cells

We adapted protocols from previously published reports to dissociate and collect whole-cell suspension from the liver. While the animal was anesthetized through continuous inhalation of 2% isoflurane, a 5 cm long incision was made in the abdomen to expose the portal vein and inferior vena cava. To perfuse the liver, the portal vein was cannulated, and ~30 ml of solution I (1x HBBS without Ca^2+^/Mg^2+^ with 1 mM EDTA, 37 °C) was passed followed by ~40-50 ml of solution II (HBSS with 0.54 uM CaCl2, 40 ug/ml Trypsin Inhibitor, 15 mM HEPES PH 7.4 and 6000 units of collagenase IV, 37 °C). The liver was excised and massaged in washing solution (DMEM + Ham’s F12 (50:50) with 5% FBS and 100 U/ml Penicillin/Streptomycin, 4 °C) to release cells from the liver capsule. The cell suspension was then passed through 40 um filter to remove doublets/undigested tissue chunks, pelleted by centrifugation at 350xg for 4 min at 4 °C to remove debris and resuspended in 15 ml wash buffer. Cells were counted with an automated hemocytometer, and ~1-1.5 million cells were pelleted and processed for library preparation.

### Dead cell removal, single cell library preparation and sequencing

MACS Dead Cell Removal Kit (Miltenyl Biotec) was used according to the manufacturer’s protocol to remove dead cells and obtain unbiased single-cell suspensions of liver cells with high viability. Following this, single-cell sequencing libraries were prepared individually from each time point using the 10X Genomics Chromium Single Cell 3’ Kit v3 and sequenced with Illumina NovaSeq 6000 on a SP/S4 flow cell to obtain 150bp paired reads.

### Raw sequencing data processing and cell-type identification

Single-cell libraries produced over four billion reads. We used Cell Ranger v3.1 pipelines from 10X Genomics to align reads and produce feature barcode matrices. Seurat v3.1 (Butler et al., 2018) was used for QC and analysis of individual feature barcode matrices were further integrated after removing batch-specific effects using BEER v0.1.7 (Zhang et al., 2019). Data was log-normalized, scaled, and clustered after PCA analysis. Hepatocyte and NPC clusters were identified based on the expression of various known marker genes. To identify cell types within the subset of NPCs, they were further subjected to unsupervised UMAP clustering.

### Pseudo-temporal trajectory analysis

Monocle v2.0 was used to perform pseudotime analysis, according to the online documentation (Qiu et al., 2017b; 2017a; Trapnell et al., 2014). The CellDataSet class monocle objects were made from log-normalized Seurat object containing the cells under consideration. Dimensionality reduction was performed using the DDRTree algorithm, and the 3000 most significant deferentially expressed genes were used as ordering genes to perform pseudotime ordering, to obtain cell trajectories. Genes with expression patterns co-varying with pseudotime were determined by the ‘differentialGeneTest()’ module, clustered and plotted using the ‘plot_pseudotime_heatmap()’ module. Expression patterns in clusters were distinguished as upregulated/downregulated along the pseudotime, and gene ontology analysis was performed using DAVID 6.8 (Dennis et al., 2003) to identify biological pathways that are up or downregulated along the pseudotime.

### Pathway scoring

Relative scores of biological pathways were assessed with the CellCycleScoring() module in Seurat v3.1. Pathway terms and exhaustive lists of candidate genes for each pathway were obtained from the Rat Genome Database (RGD) (Smith et al., 2020). The summary heatmap was generated using Seurat v3.1, considering all pathways with at least 50 genes. Box plots to demonstrate cell cycle and pathway scores were constructed using ggplot2, and pairwise comparison with reference using a T-test was performed with the ggpubr package.

### Gene regulatory network analysis

SCENIC pipeline (Aibar et al., 2017) was used to estimate the AUCell Score activity matrix from the log-normalized Seurat object containing the subset of hepatocytes. Unlike the standard SCENIC workflow where this AUCell score activity matrix is binarized by thresholding to generate binary regulon-activity matrix, we retained the full AUCell score for all further analysis. UMAP plots, heatmaps and violin plots demonstrating regulon activities based on AUCell scores were made in Seurat 3.1. AUCell scores were plotted over pseudotime cell-trajectories using Monocle 2.0.

### Imputation and Cell-cell communication analysis

ScRNA-seq data often contains dropouts or missing values due to failure in the detection of RNAs (Kharchenko et al., 2014). In our dataset, we noticed that NPCs have lower UMIs relative to hepatocytes, and could potentially have dropouts leading to incomplete representations. Hence, imputation using the MAGIC algorithm (van Dijk et al., 2018) was performed to correct for any missing values in the NPC dataset before interpreting cell-cell interactions. We constructed cell-cell communication networks and performed statistics of interactions using methods previously described in detail by Farbehi et al. (Farbehi et al., 2019). Briefly, we used a directed and weighted network with four layers of nodes, namely, the *source* cell populations expressing the ligands, the *ligands* expressed by the source populations, the *receptors* targeted by the ligands, and the *target* cell populations. Weights of edges that connect ‘*source* to *ligand*’ and ‘*receptors* to *target*’ were computed as Log2 fold change in expression of ligand/receptor in source/target compared to remaining cells. Ligand-receptor interactions were determined using a mouse-specific ligand-receptor interaction dataset compiled previously ((Farbehi et al., 2019). The sum of weights along the path was used to calculate path weights connecting a source to target through a ligand:receptor interaction. Cells were grouped according to cell types, and hepatocytes were subdivided based on cellular states. We used all ligand:receptor connections with a minimum pathweight of 1.5 and calculated the overall weight, *ws:t*, as the sum of all pathweights between the corresponding source and target. Only edges with Benjamini-Hochberg adjusted P-values, Pw<0.01 were considered significant. Further, we constructed ligand:receptor interaction dot plots using ggPlot2.

## Supporting information

Supplementary data file 1 P14__Ligand_receptor_interactions

Supplementary data file 2 Adult_Ligand_receptor_interactions

Supplementary data file 3 PHx24_Ligand_receptor_interactions

Supplementary data file 4 PHx48_Ligand_receptor_interactions

Supplementary data file 5 PHx96__Ligand_receptor_interactions

## DATA AVAILABILITY

All raw RNA-seq data files are available for download from NCBI Gene Expression Omnibus (http://www.ncbi.nlm.nih.gov/geo/) under accession number GSE151309.

## ACKNOWLEDGEMENTS

Work in the Kalsotra laboratory is supported by National Institute of Health (R01HL126845, R01AA010154), Muscular Dystrophy Association (MDA514335), Planning Grant Award from the Cancer Center @ Illinois, and the Beckman Fellowship from the Center for Advanced Study at the University of Illinois Urbana-Champaign. U.V.C. is supported by the Herbert E. Carter fellowship in Biochemistry, UIUC. S.B is supported by the NIH Tissue microenvironment training program (T32-EB019944). We acknowledge support from the Transgenic mouse core, High-throughput sequencing and genotyping core and Histology-microscopy core facilities.

## AUTHOR CONTRIBUTIONS

U.V.C., S.B. and A.K. conceived the project and designed the experiments. U.V.C. and S.B. performed experiments and analyzed the data. M.H. provided guidance with bioinformatics analyses. U.V.C., S.B. and A.K. interpreted results and wrote the manuscript. All authors discussed the results and edited the manuscript.

## COMPETING INTERESTS

The authors declare no competing financial interests.

Requests for materials should be sent to A.K. at

---

**Figure 1—figure supplement 1.**
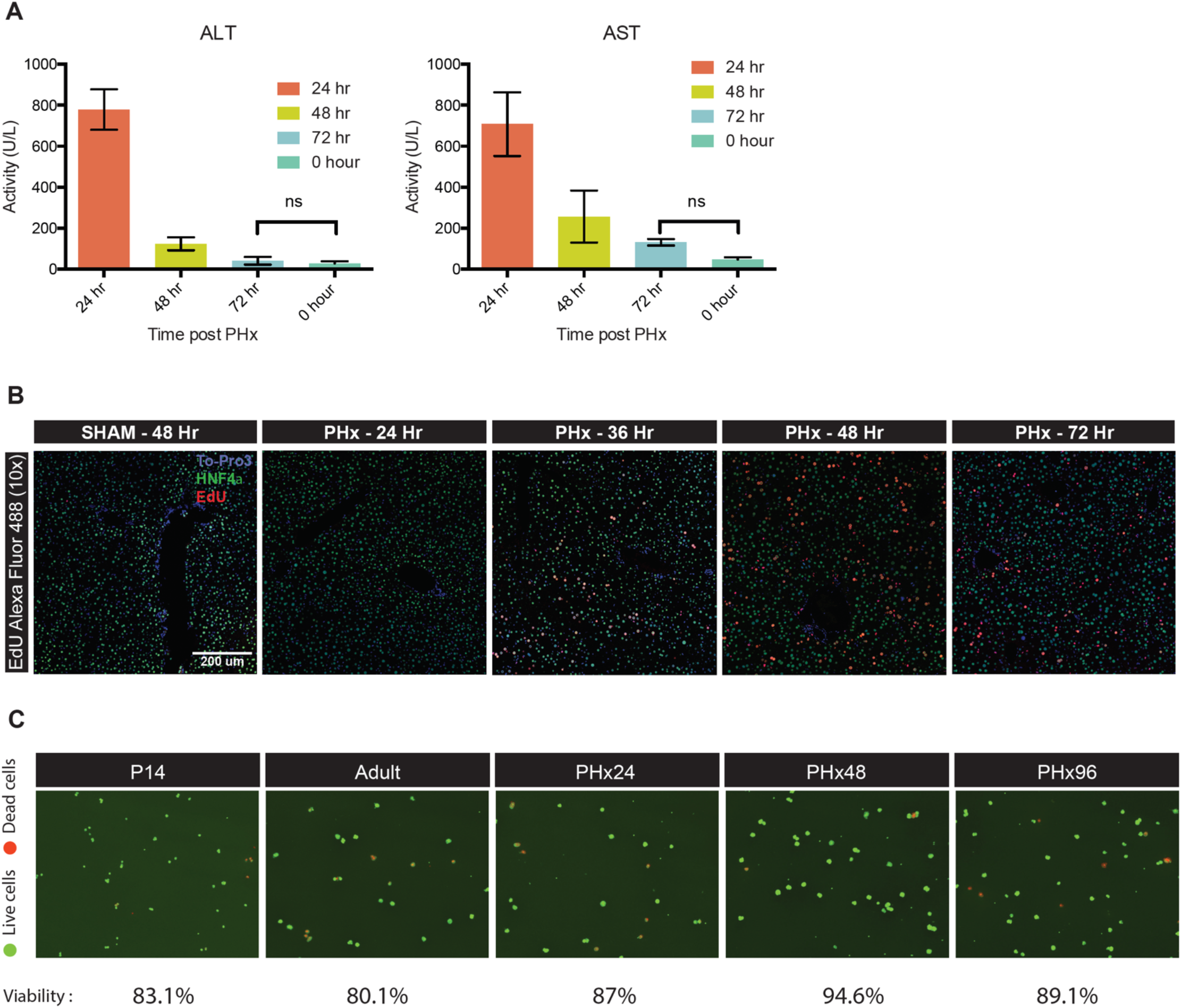
**(A)** ALT and AST levels 24 hours, 48 hours, 72 hours post-PHx and 0 hr(control SHAM mice). Levels of serum injury markers (ALT and AST) are restored to normal levels within 72 hours after PHx. Data are mean ± s.d. *P < 0.05, ns: not significant, (n = 3 animals per group) **(B)** Fluorescent imaging of hepatocyte proliferation measured by in vivo EdU incorporation in post-PHx and SHAM livers. White arrows indicate proliferating hepatocytes (Co-labeled for Hnf4α in green, incorporated EdU in red, and nuclei in blue). Images taken under 10X resolution are shown. **(C)** Fluorescent imaging of hepatocytes indicating the viability of cells used for scRNA-seq. Dead/dying cells are stained in red (Propidium Iodide) and viable cells are stained in green (Acridine Orange). Percentage viability was calculated by Nexcelom Cellometer is also shown.

**Figure 1—figure supplement 2.**
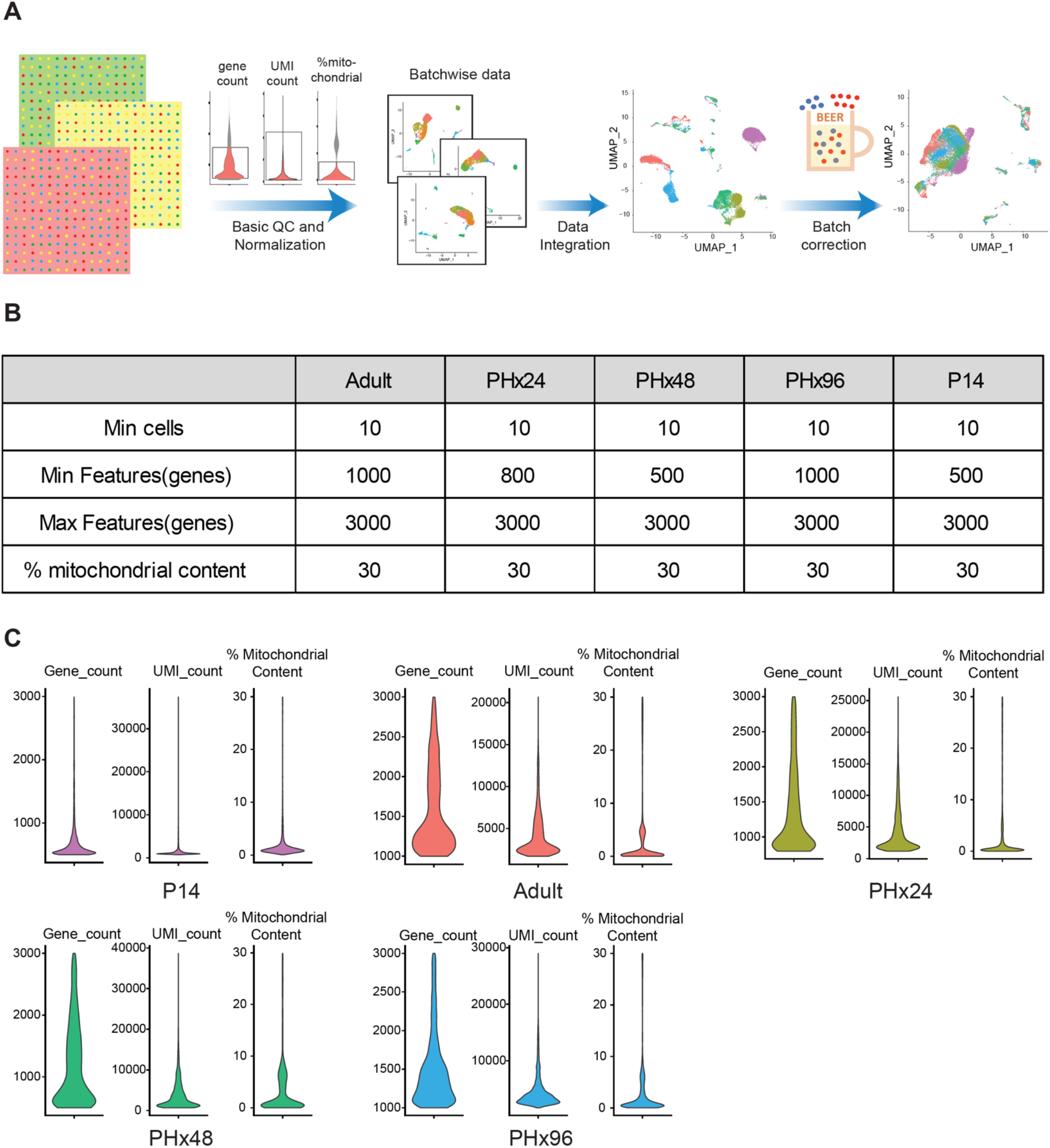
**(A)** Computational workflow depicting data processing and analysis pipeline for scRNA-seq data. Cell Ranger was used to align raw reads and generate feature-barcode matrices from scRNA-seq output for each sample. Seurat v3.1 was used to perform basic QC (see suppl. Table) and normalization, after which data was integrated using BEER to remove extraneous batch specific effects. **(B)** Table showing Quality Check (QC) cutoffs applied on raw reads from scRNA-seq data prior to the analysis. **(C)** Violin plots showing the distribution of gene counts, UMI counts and % mitochondrial content in P14, Adult, PHx24, PHx48 and PHx96 samples, after applying QC cutoffs listed above to scRNA-seq data. These set of cells were used for all downstream analysis.

**Figure 1—figure supplement 3.**
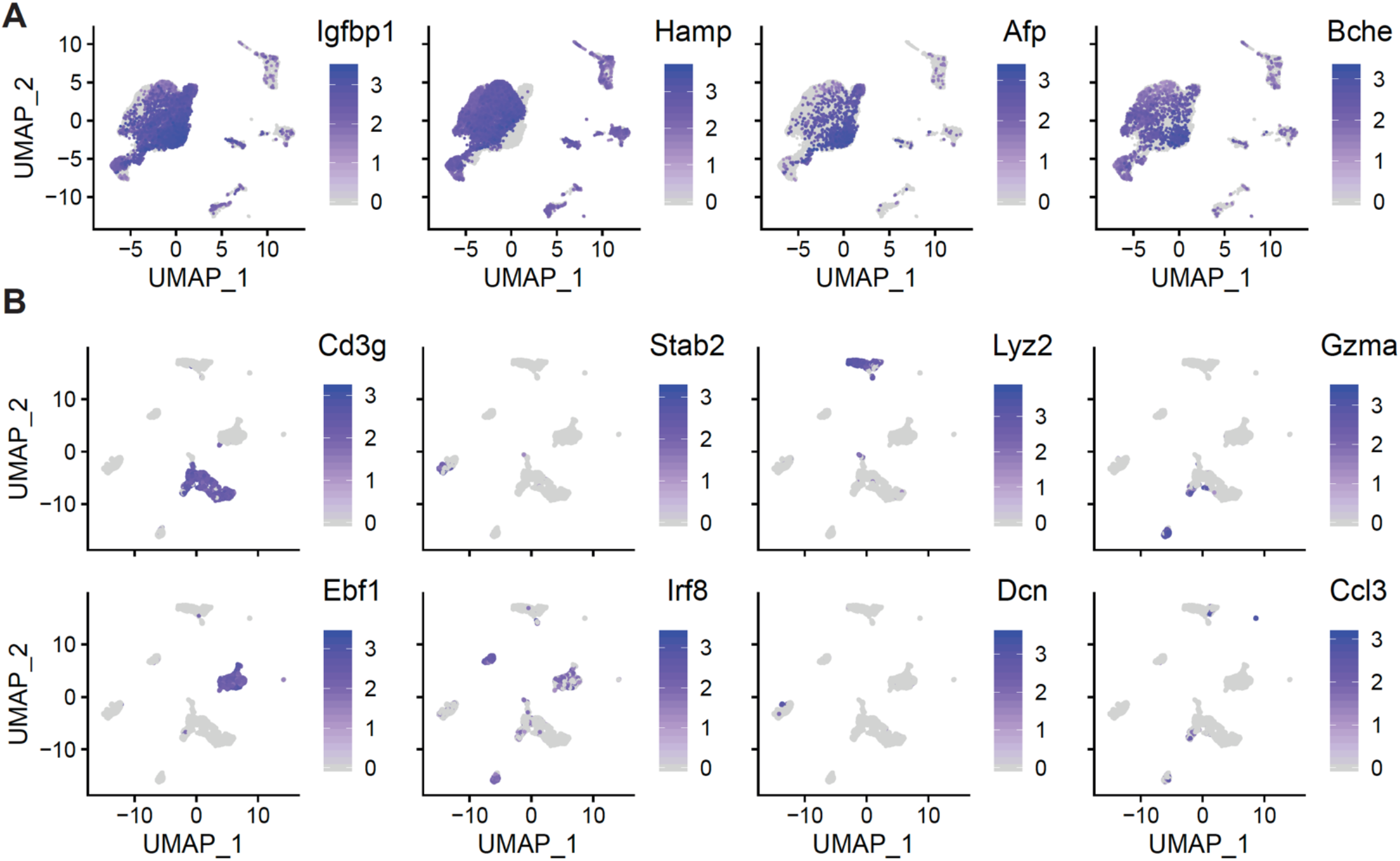
**(A)** UMAP plots representing the expression of hepatocyte-specific genes. **(B)** NPC subpopulation UMAP plots representing the expression of genes specific to different NPC populations identified. Cells in (A) and (B) are colored by the expression levels of the indicated gene, as calculated with Seurat V3.1.

**Figure 1—figure supplement 4.**
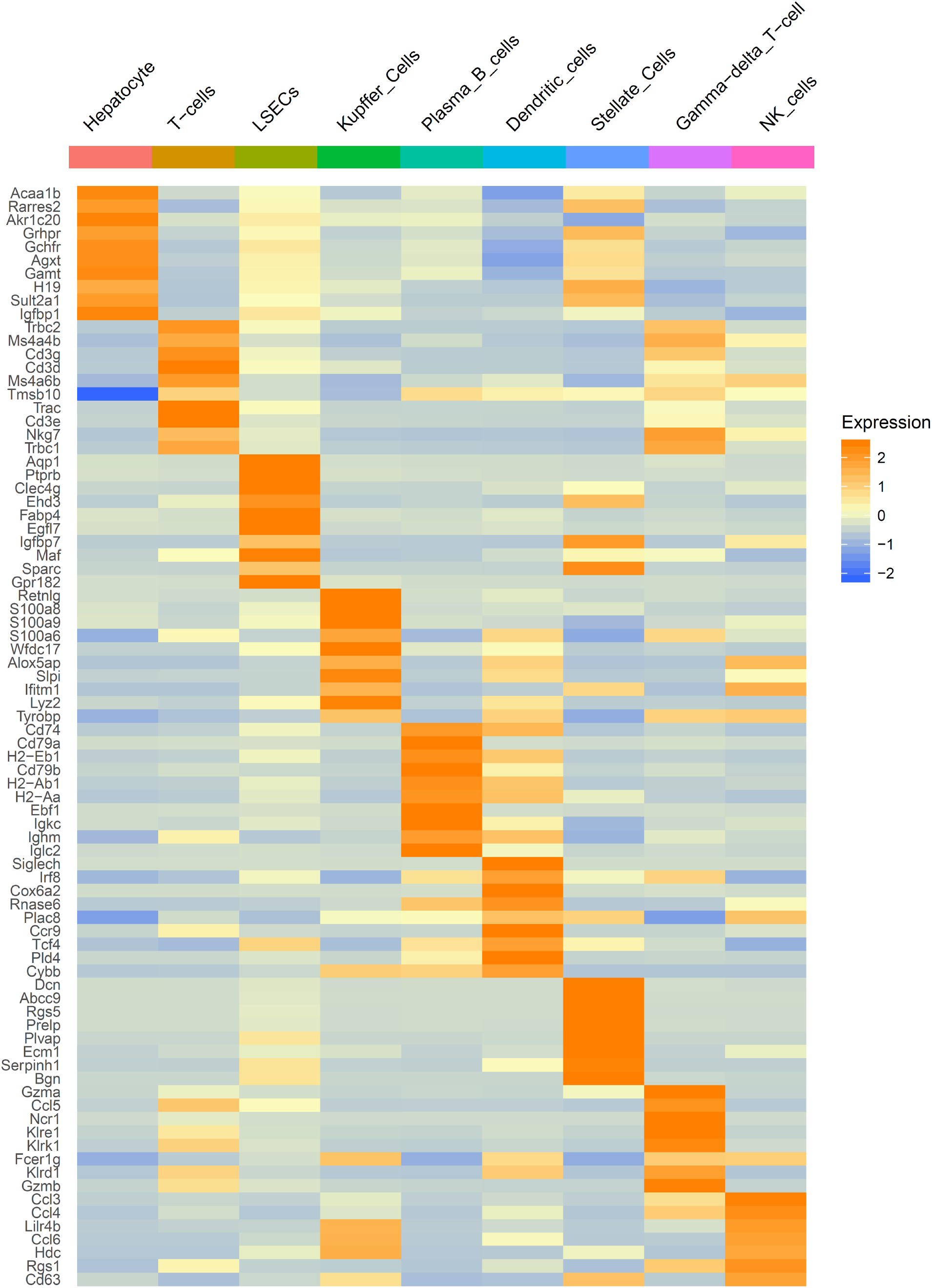
Heatmap showing top genes that were enriched (>2 fold) in each cell type cluster. Scale bar shows relative gene expression.

**Figure 3—figure supplement 1.**
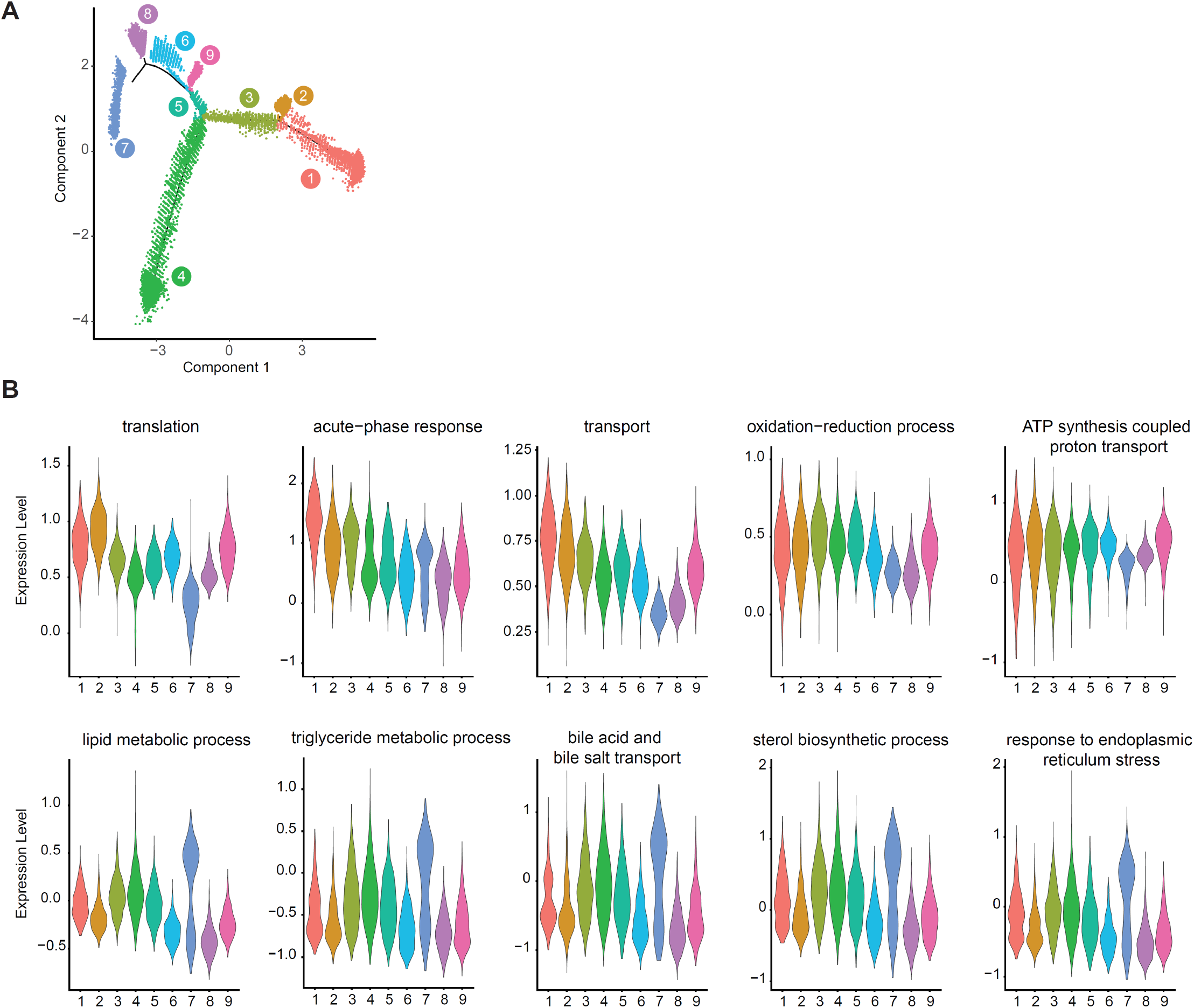
**(A)** DDRTree trajectory showing the nine states identified by Monocle2 along the pseudotime trajectory for hepatocytes from all timepoints. **(B)** Violin plot showing relative scoring (using Seurat3.1) for top ontology categories differentially regulated (with respect to the trajectory bifurcation) during regeneration for hepatocytes belonging to states identified in **A**.

**Figure 4—figure supplement 1.**
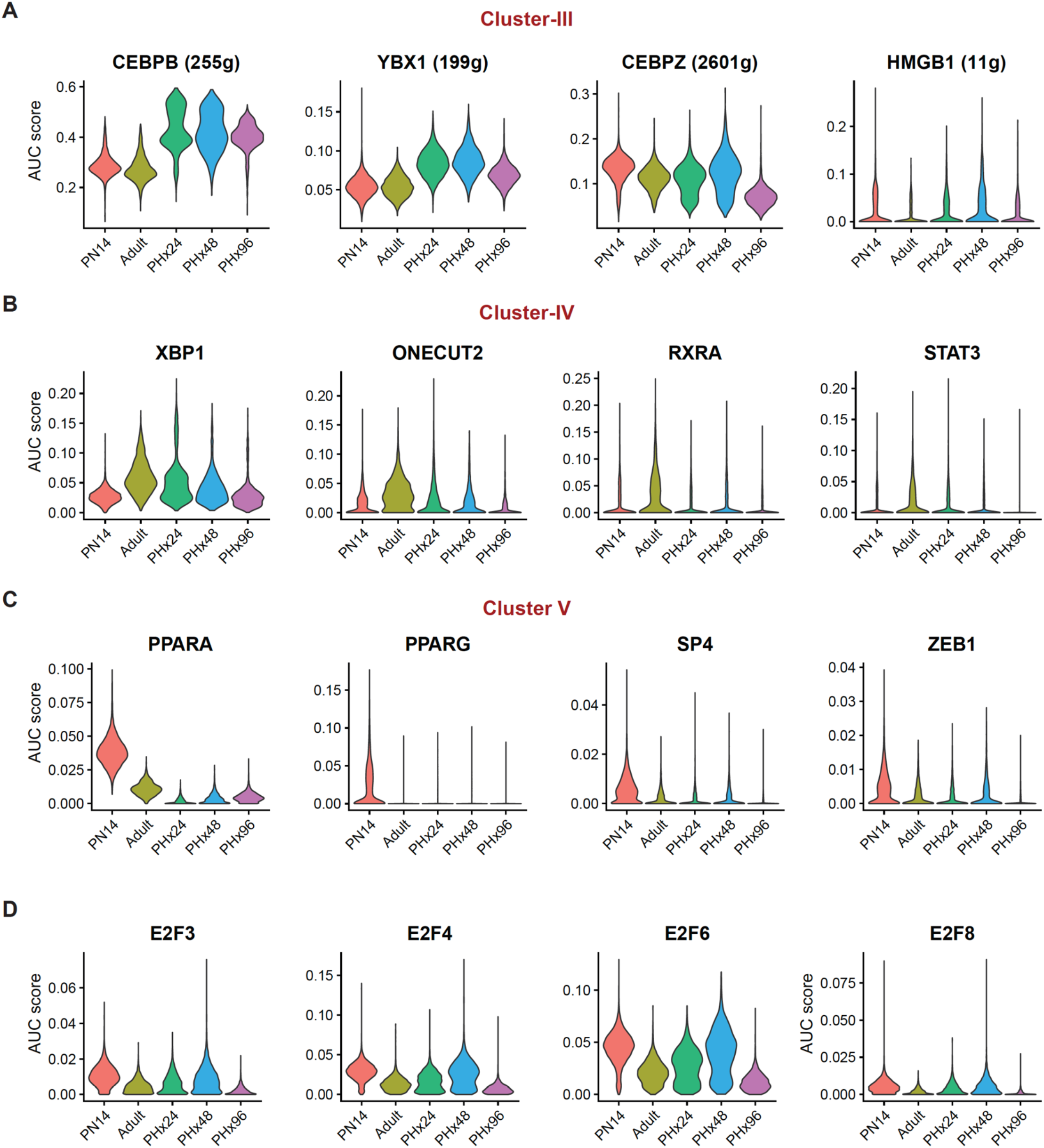
Violin plots showing AUC-score distribution within hepatocytes at different time points, for the regulons belonging to **(A)** Cluster III **(B)** Cluster IV **(C)** Cluster V, and **(D)** various E2F transcription factors.

**Figure 4—figure supplement 2.**
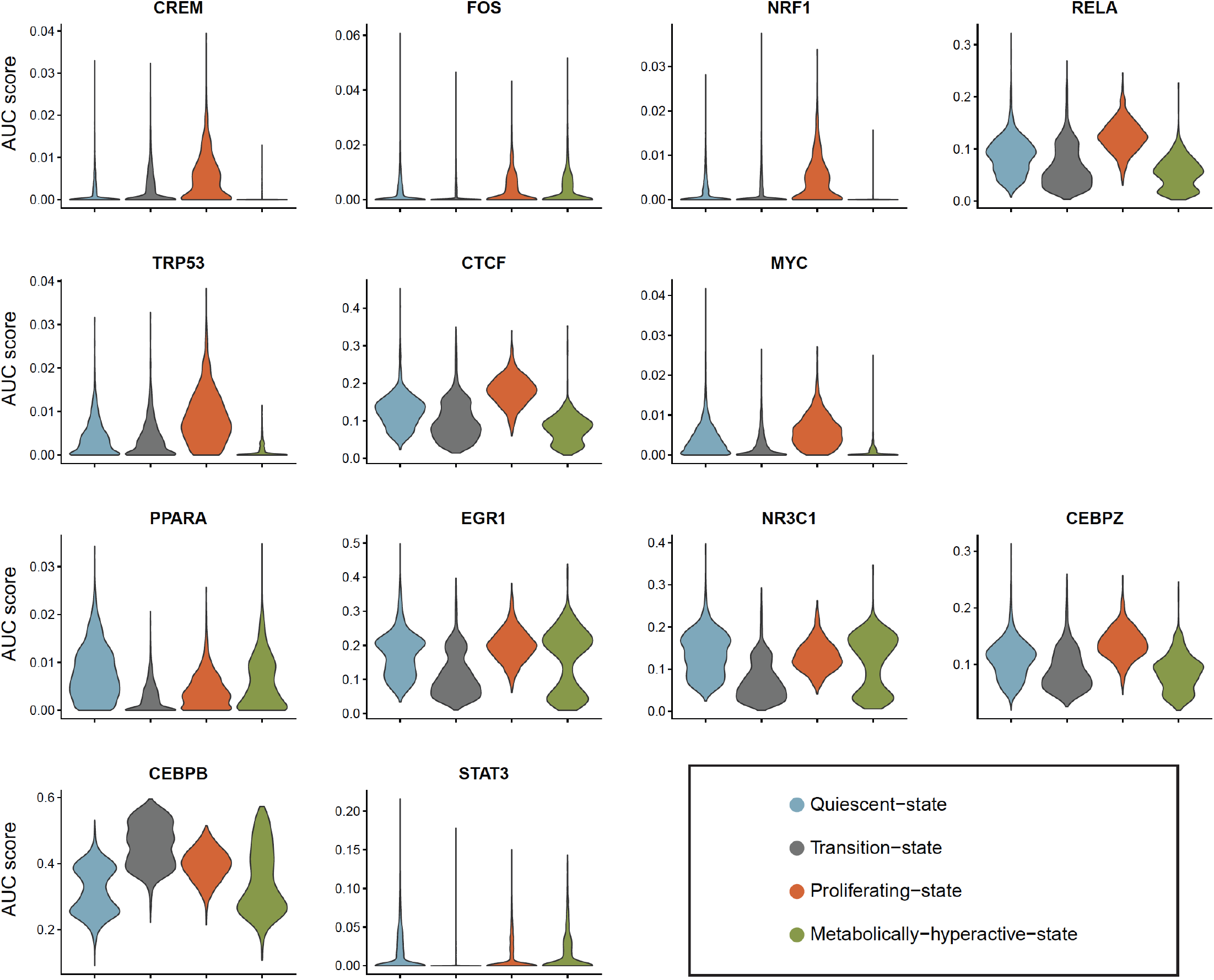
Violin plots showing state-wise distribution of regulon AUC-scores within hepatocytes.

**Figure 5—figure supplement 1.**
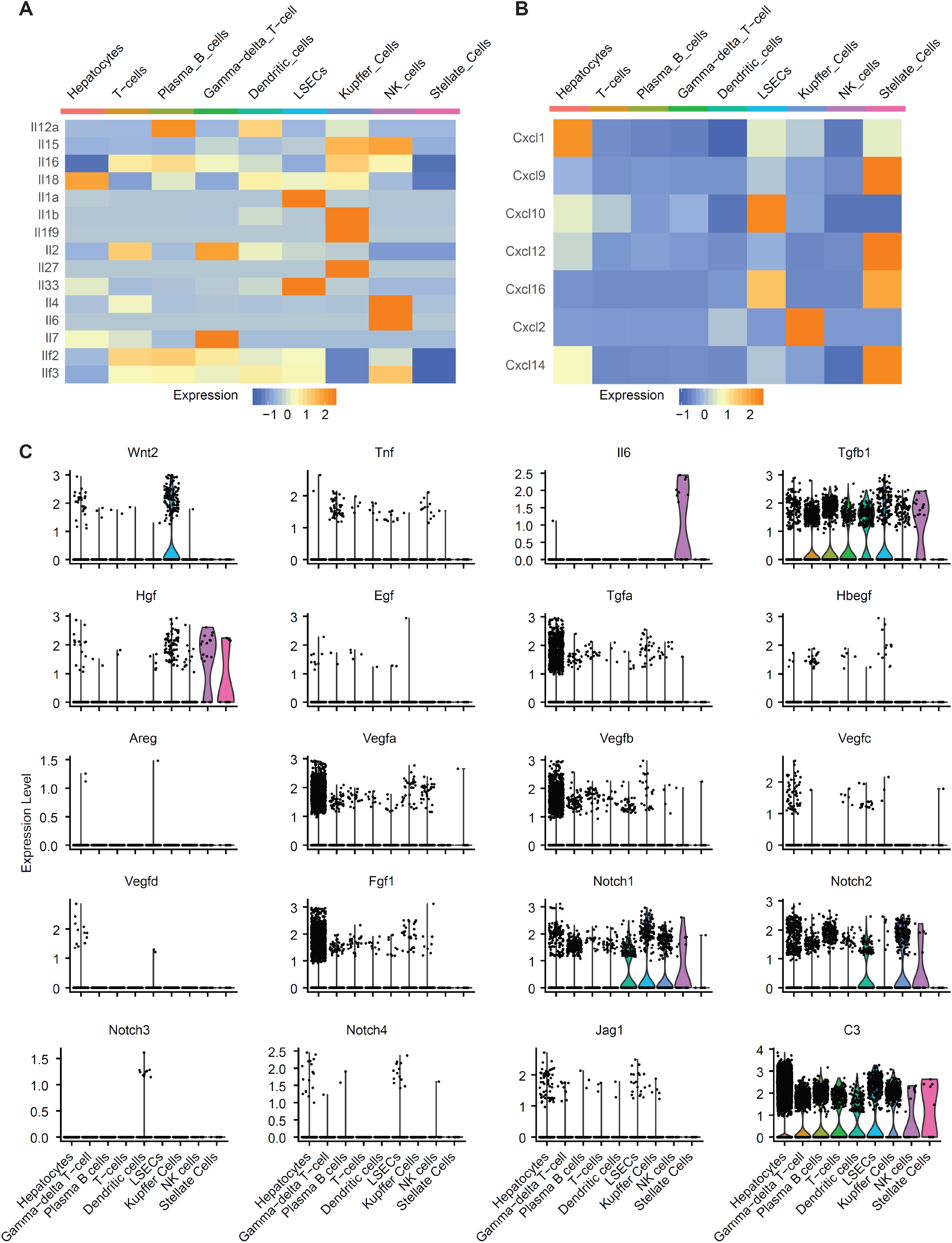
Heatmap demonstrating cell-type-specific expression patterns of **(A)** Interleukins and **(B)** Chemokines identified in our dataset. **(C)** Violin Plots showing cell-type-specific expression patterns of different mitogens identified in our dataset.

**Figure 5—figure supplement 2.**
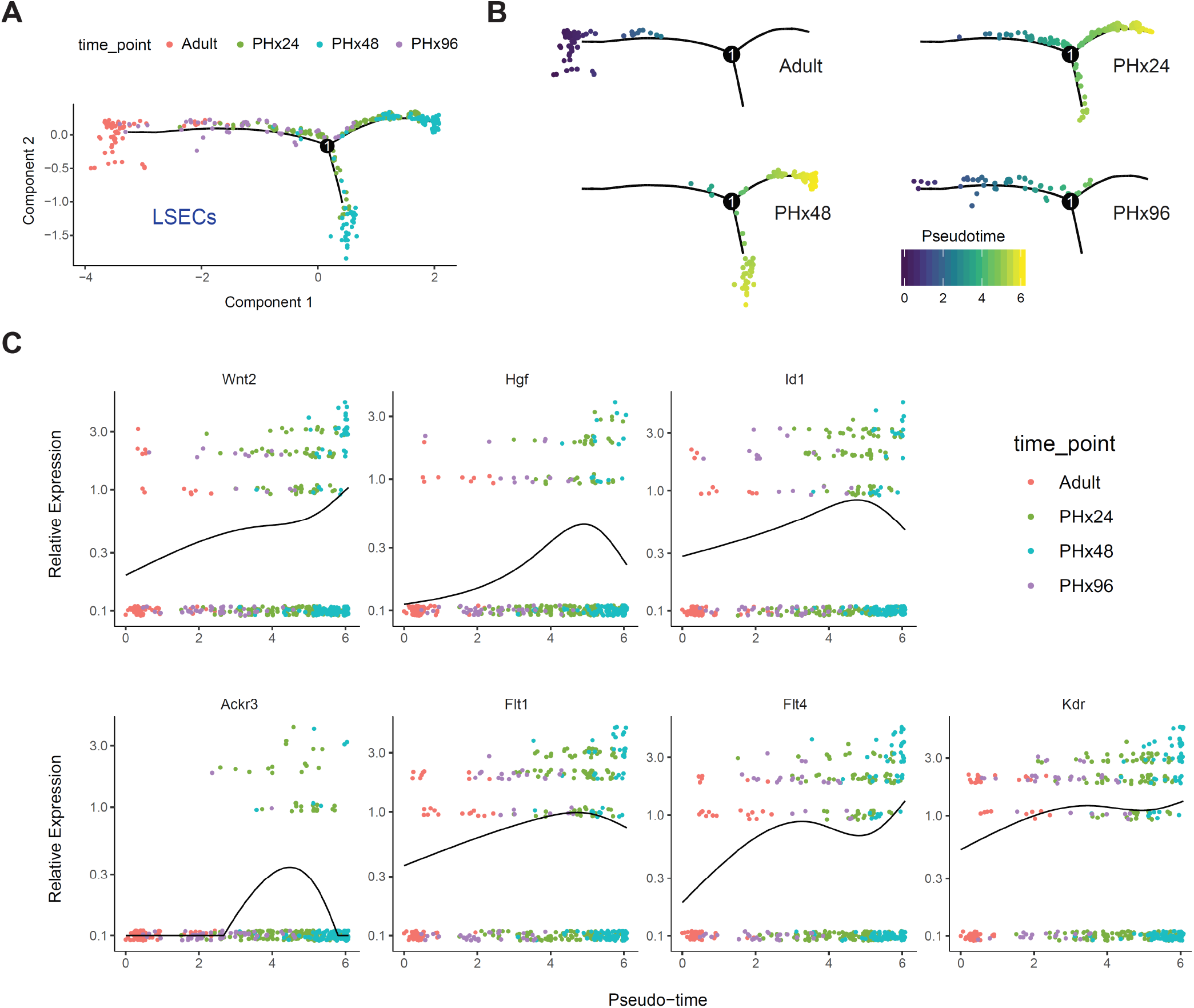
**(A)** LSEC activation during regeneration. Single cell trajectories were constructed and pseudotime values were calculated using Monocle 2, with LSECs identified from adult, PHx24, PHx48 and PHx96 samples. Cells within trajectories are colored by sample identity. **(B)** Distribution of LSECs from different time points along the activation trajectory. Cells are colored by pseudotime value. **(C)** Variation in relative expression levels of LSEC activation-associated genes along the pseudotime trajectory. X- and Y-axis corresponds to pseudo-time value of each cell and relative expression level of the gene under evaluation respectively.

**Figure 5—figure supplement 3.**
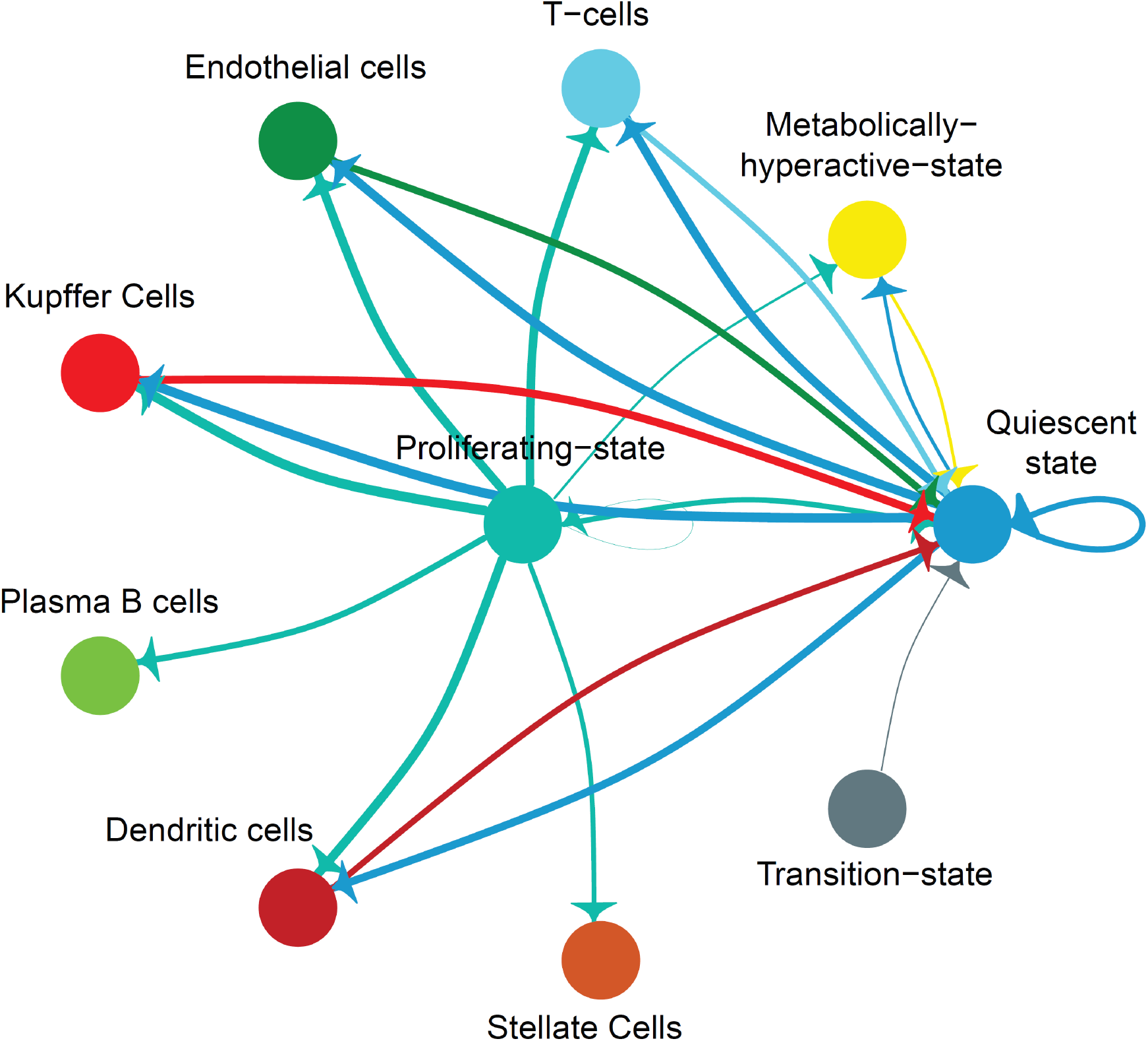
Network diagram showing significant cell-cell interactions for P14, where interactions are indicated by arrows(edges) pointing in the source to target direction. Thickness indicates the sum of weighted paths between populations and color of arrows corresponds to the source

## Key Resource Table

| Reagent type<br>(species) or resource | Designation | Source or reference | Identifiers | Additional<br>information |
| --- | --- | --- | --- | --- |
| Strain, strain background | Mouse: C57BL/6J | The Jackson Laboratory, Bar Harbor, ME | [Stock# 00664, https://www.jax.org/strain/000664](https://www.jax.org/strain/000664) |  |
| Antibody | Anti-Hnf4a C11F12 (rabbit monoclonal) | Cell Signaling Technology | Catalog# 3113 | IF (1:500) |
| Antibody | Anti-Rabbit IgG(H+L) DyLight 594 | Thermo Fisher Scientific | Catalog # 35561 | Cross-Adsorbed Secondary Antibody(1:500) |
| Nuclear stain | TO-PRO™-3 Iodide (642/661) | Thermo Fisher Scientific | Catalog# T3605 |  |
| Commercial assay or kit | Click-iT™ EdU Cell Proliferation Kit | Thermo Fisher Scientific | Catalog# C10337 |  |
| Reagent | 5-Ethynyl-2'-deoxyuridine (EdU) | Cayman chemical company | Catalog# 20518 |  |
| Other | Fisherbrand Micro Blood Collecting Tubes | Thermo Fisher Scientific | Catalog# 02-668-10 |  |
| Other | BD Microtainer™ Capillary Blood Collector | Thermo Fisher Scientific | Catalog# 02-675-186 |  |
| Commercial assay or kit | AST/GOT Reagent | Thermo Fisher Scientific | Catalog# TR70121 |  |
| Commercial assay or kit | ALT/GPT Reagent | Thermo Fisher Scientific | Catalog# TR71121 |  |
| Other | 4/0 Silk Sutures | KW-MED | Catalog# 683S | BRAIDED, BLACK,NFS-2, 18" |
| Other | 5/0 Silk Suture | Medrep express | Catalog# 682S | FS-2, 18" |
| Other | 7 mm Reflex clips | World Precision Instruments | Part #500344 |  |
| Reagent | Collagenase, Type 4 | Worthington Biochemical Corporation | Catalog# LS004188 |  |
| Commercial kit | Dead Cell Removal Kit | Miltenyi Biotec | Order no: 130-090-101 |  |
| Commercial kit | MS Columns | Miltenyi Biotec | Order no: 130-042-201 |  |
| Software/algorithm | CellRanger | 10 X Genomics | https://support.10xgenomics.com/single-cell-gene-expression/software/downloads/latest |  |
| Software/algorithm | Seurat v 3.1.0 | PMID: 29608179 | https://satijalab.org/seurat/ |  |
| Software/algorithm | BEER v 0.1.7 | PMID: 31636959 | https://github.com/jumphone/BEER |  |
| Software/algorithm | Monocle v 2.0 | PMID: 24658644 | http://cole-trapnell-lab.github.io/monocle-release/docs/ |  |
| Software/algorithm | SCENIC | PMID: 28991892 | https://github.com/aertslab/SCENIC |  |
| Software/algorithm | Cell communication analysis | PMID: 30912746 | https://elifesciences.org/articles/43882/figures#code1 |  |

