## Supplementary figures and images for "Cellular plasticity balances the metabolic and proliferation dynamics of a regenerating liver"

### Supplementary data file 1 P14__Ligand_receptor_interactions

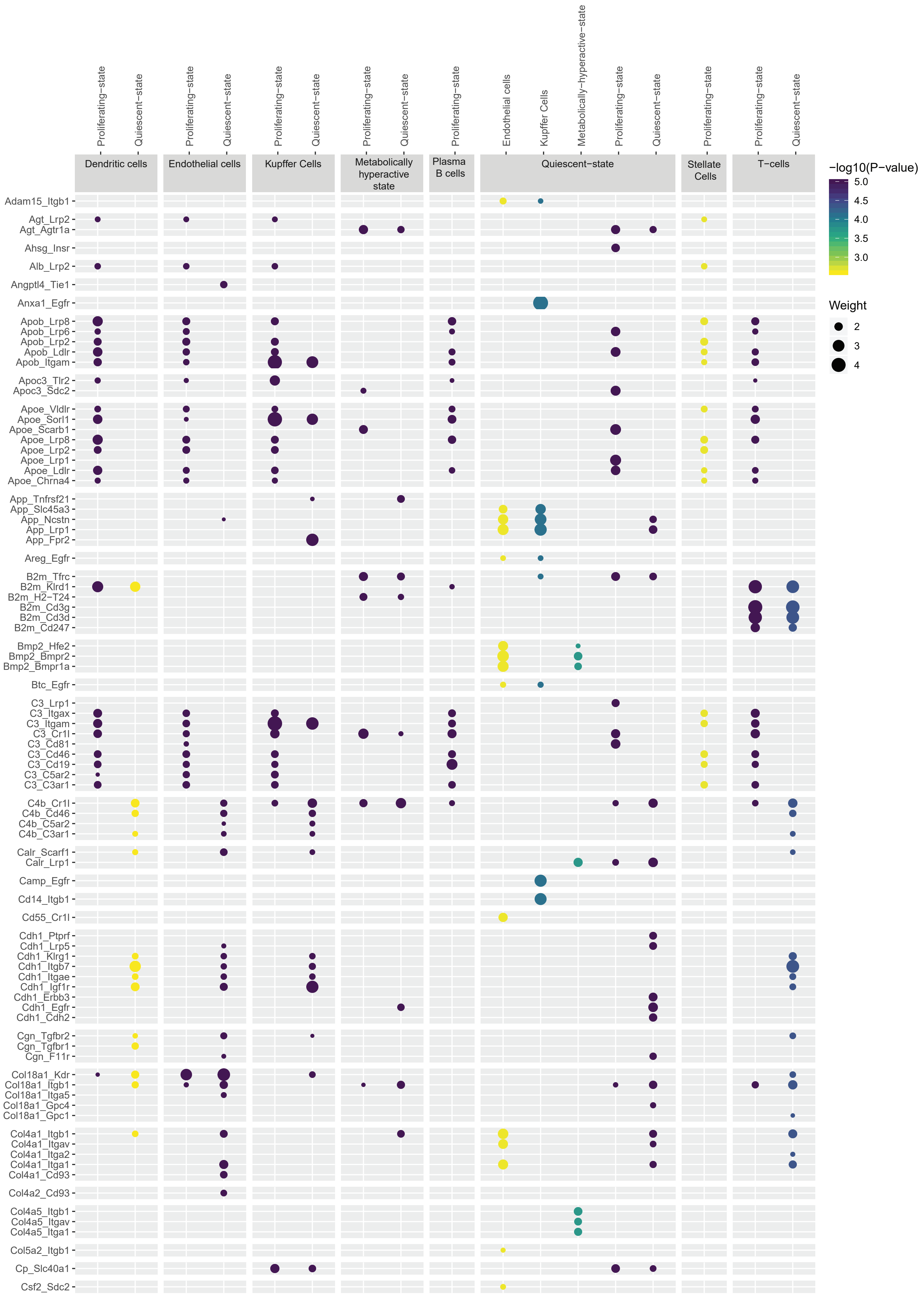

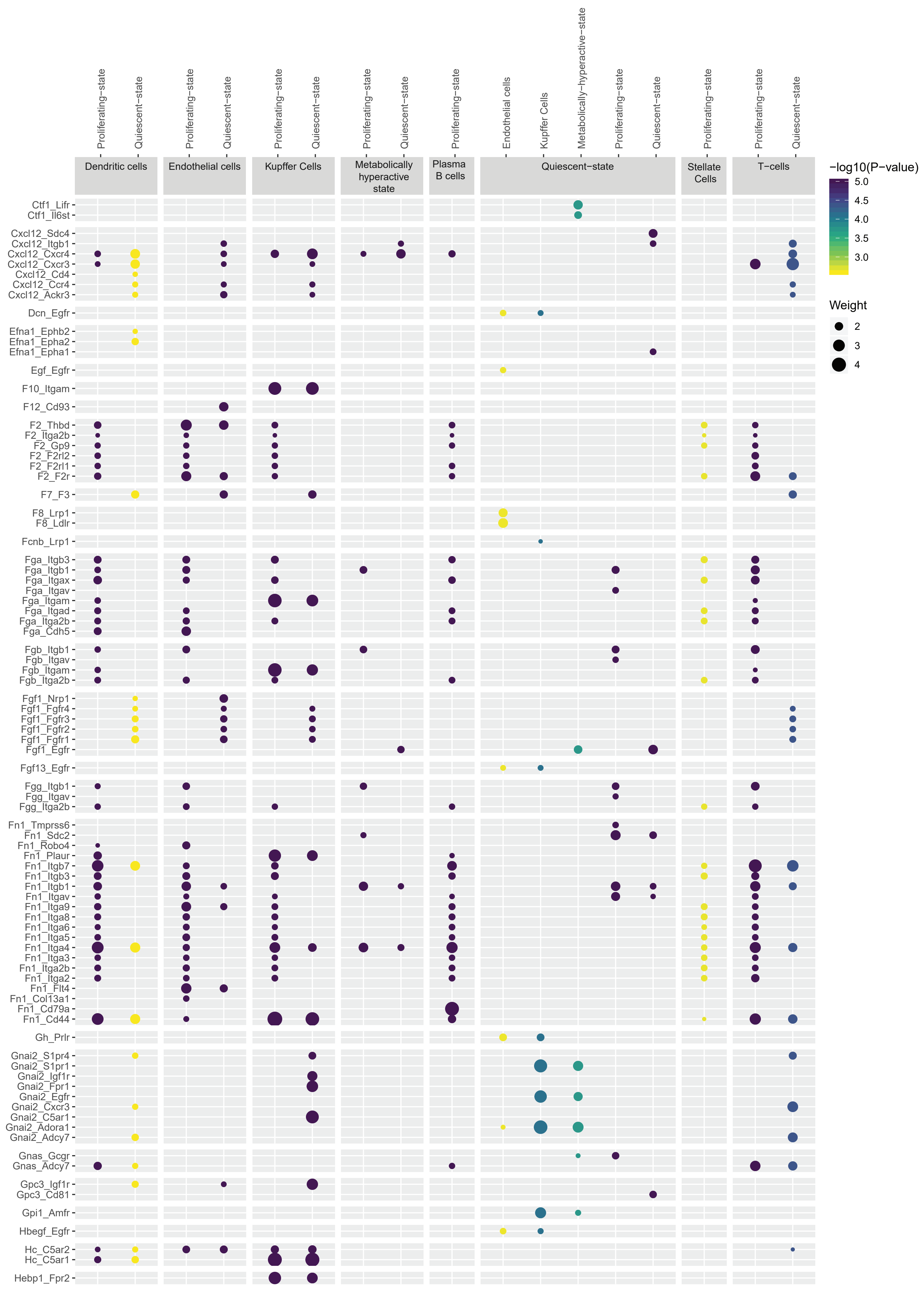

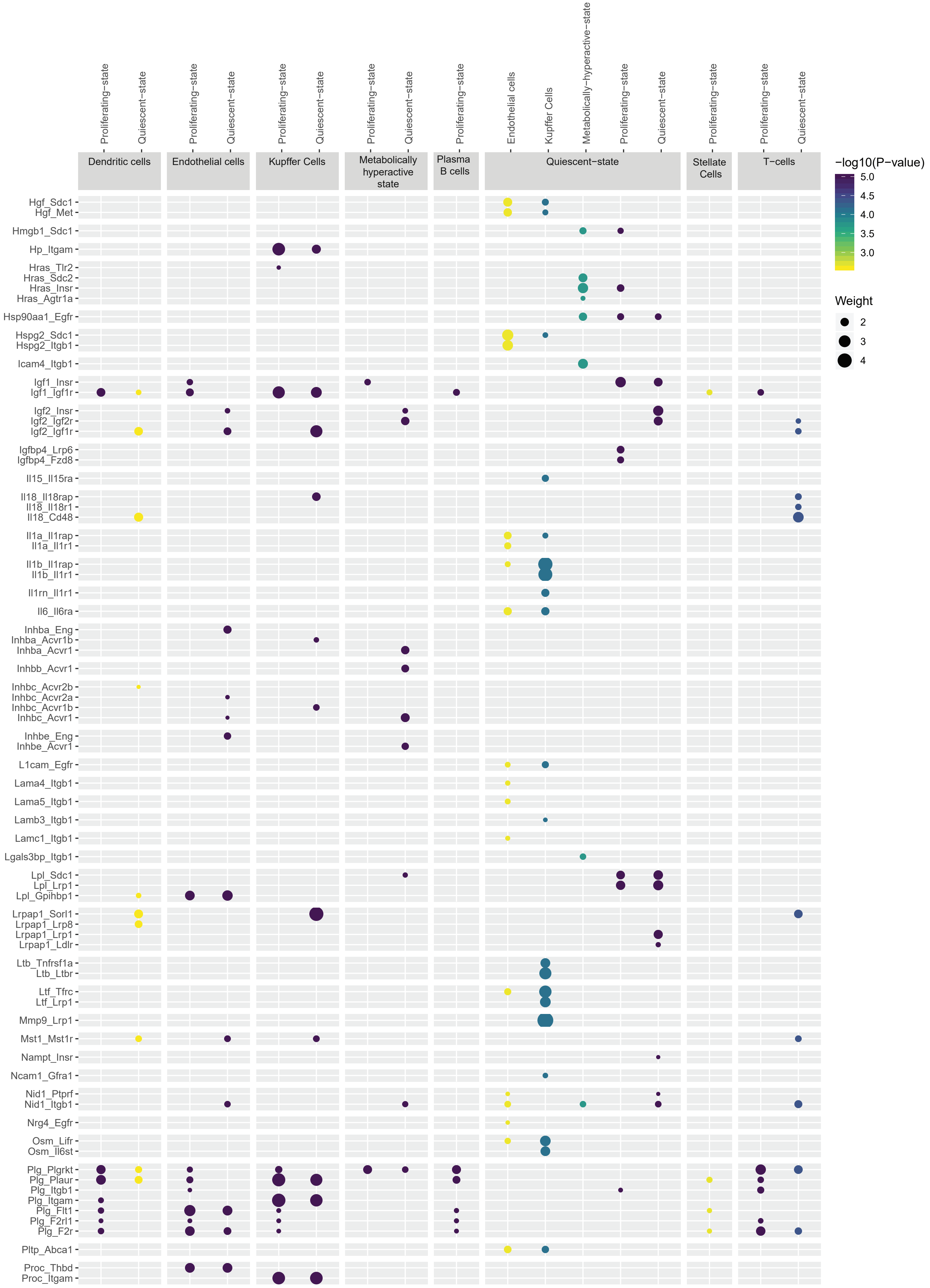

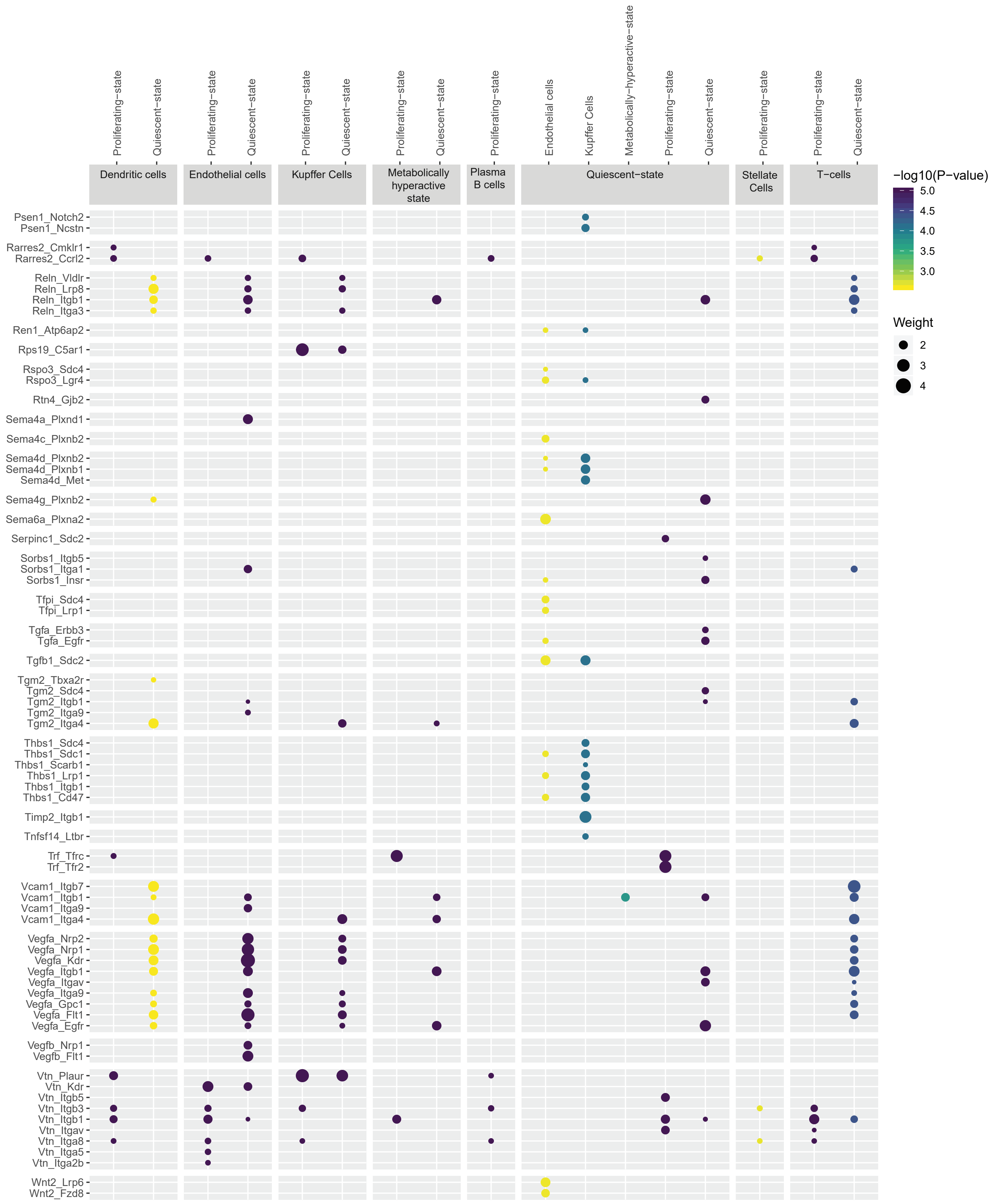

### Supplementary data file 2 Adult_Ligand_receptor_interactions

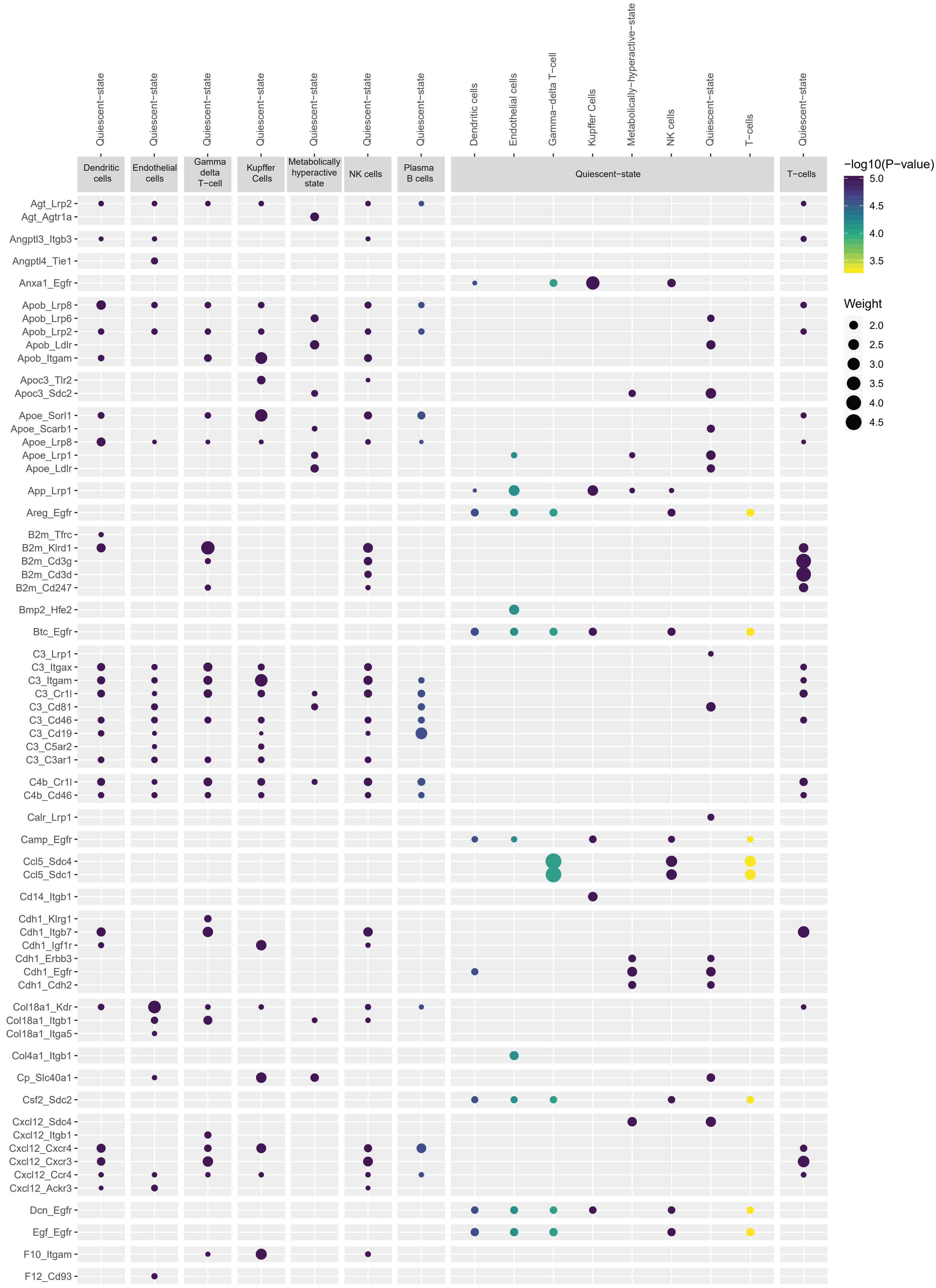

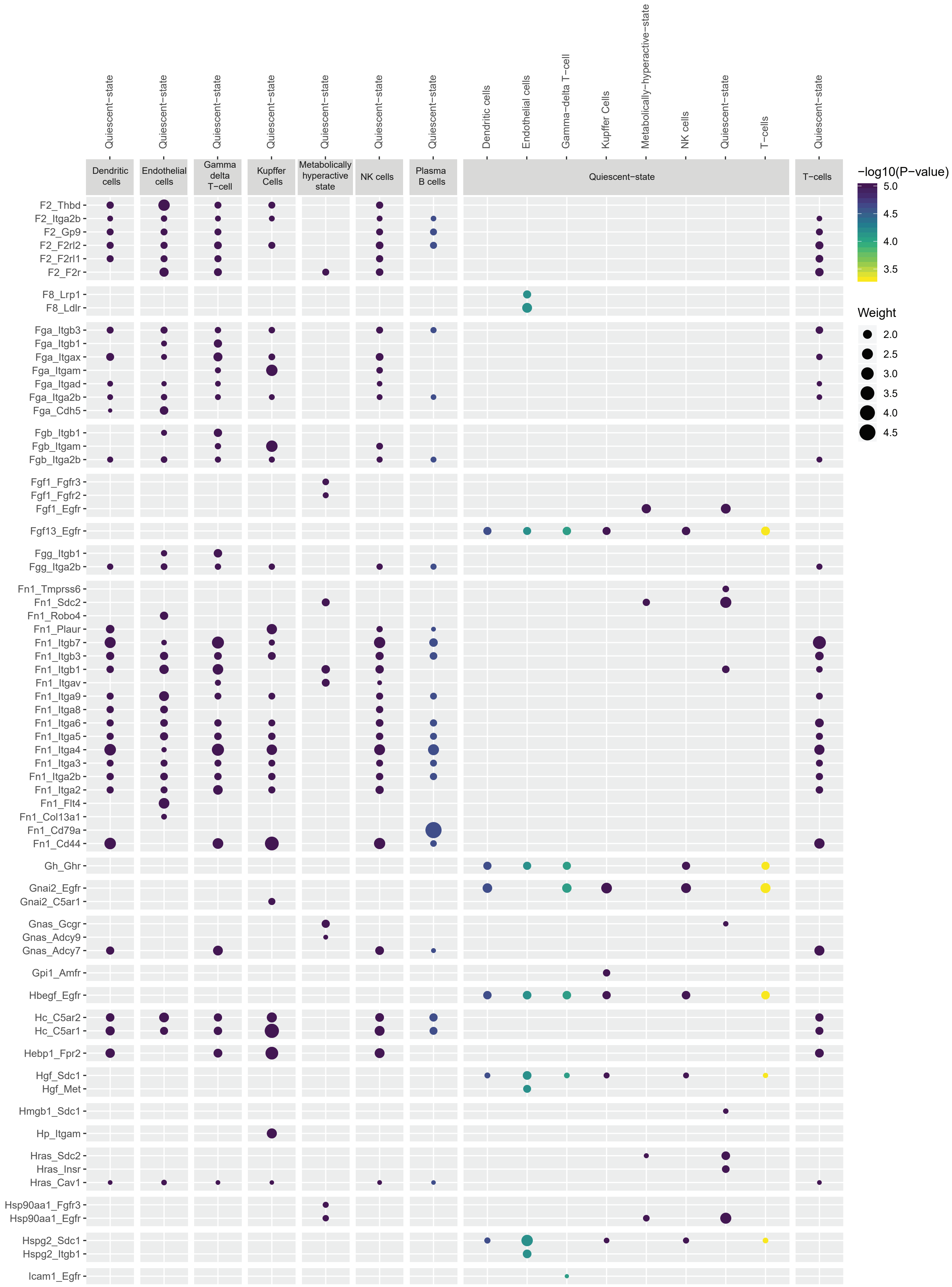

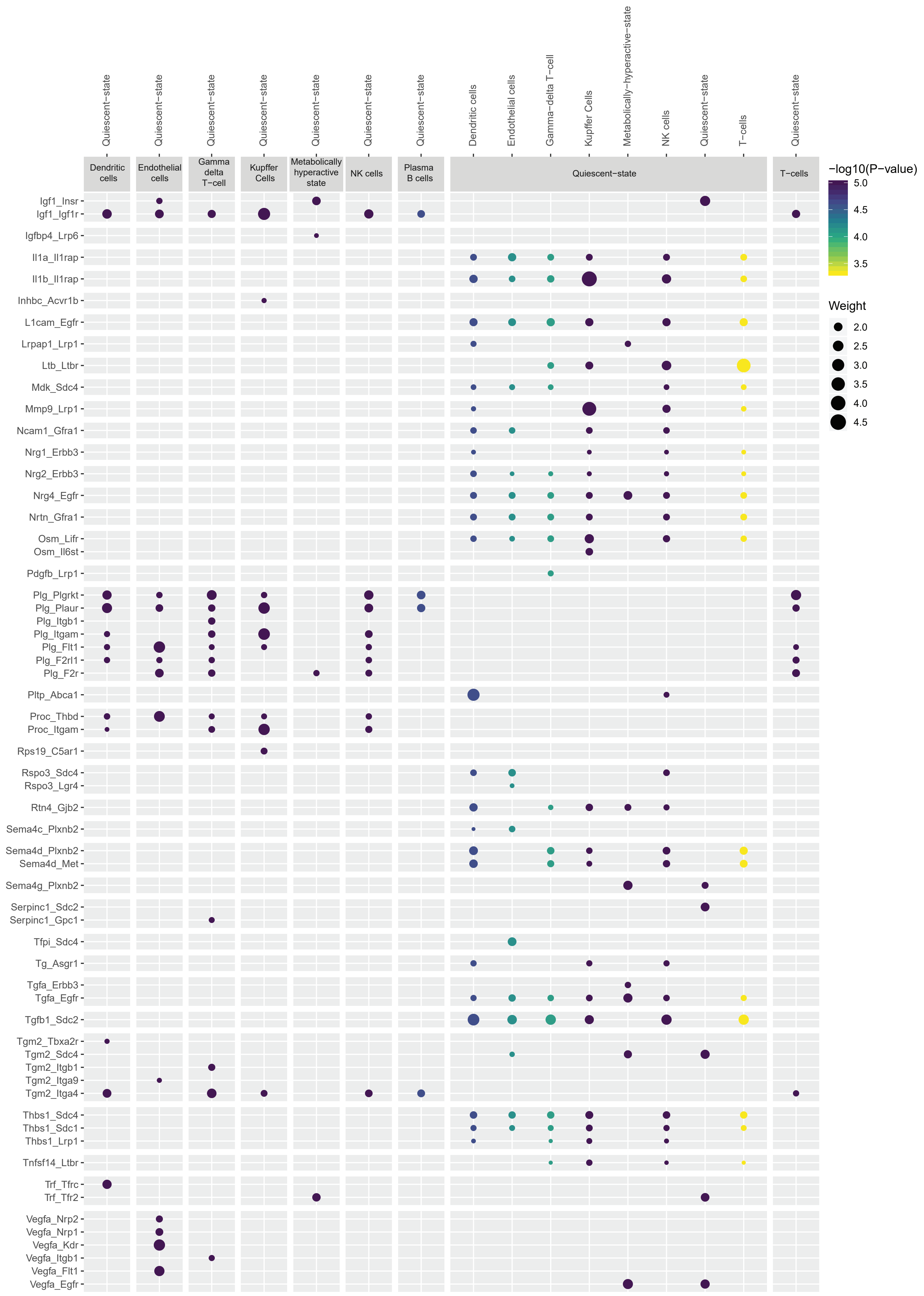

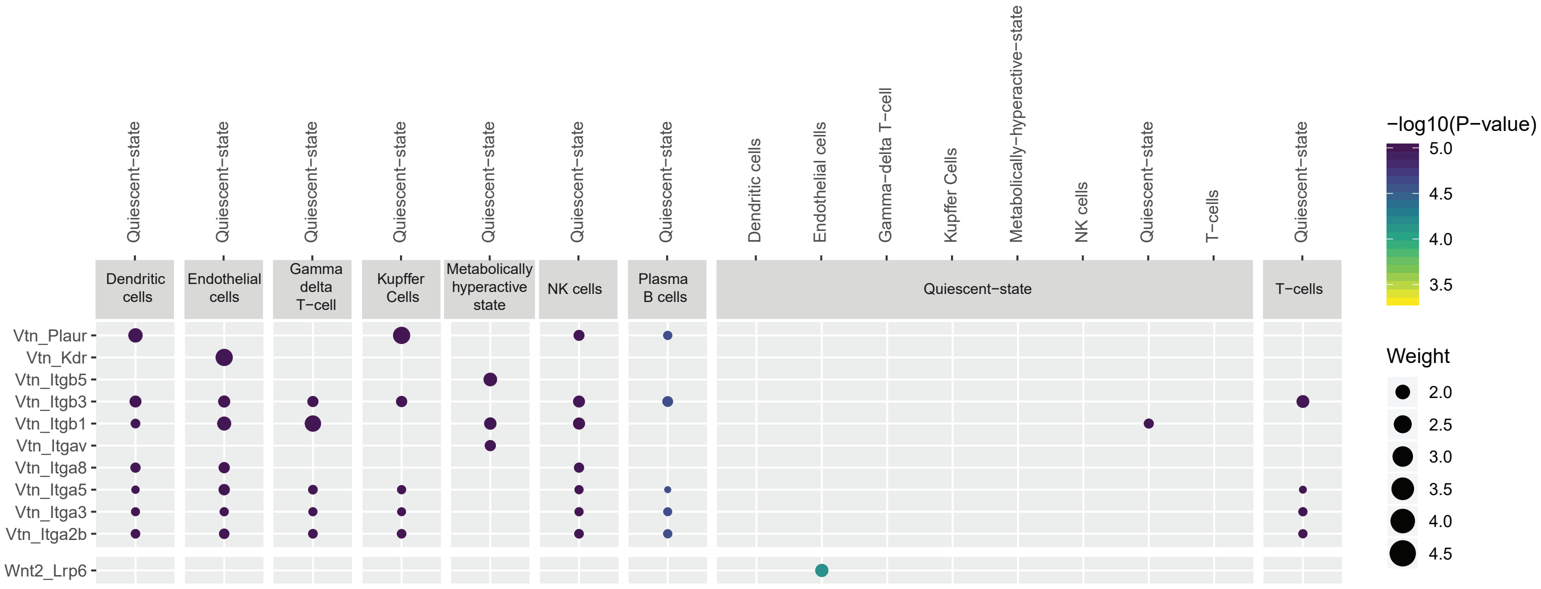

### Supplementary data file 3 PHx24_Ligand_receptor_interactions

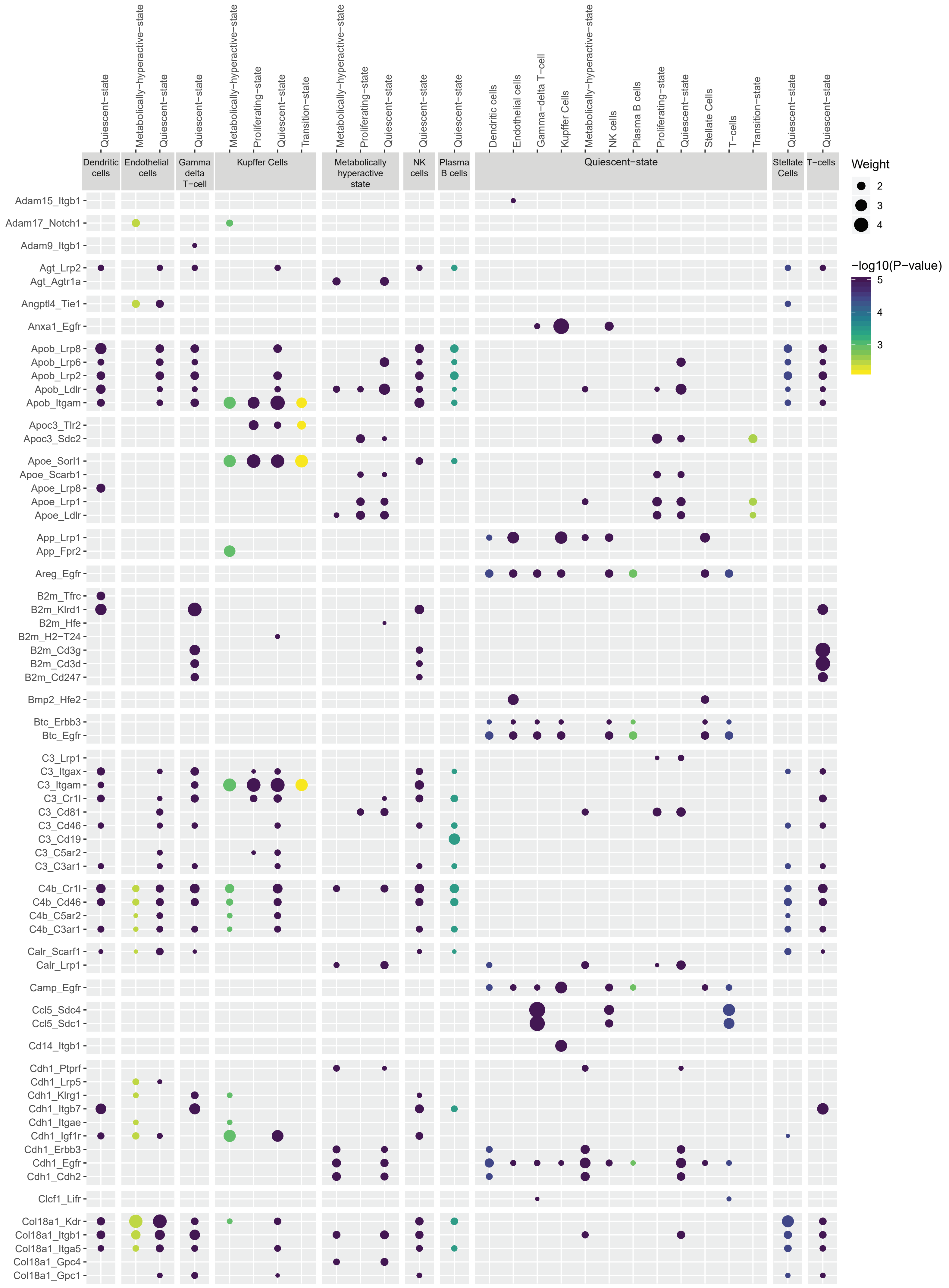

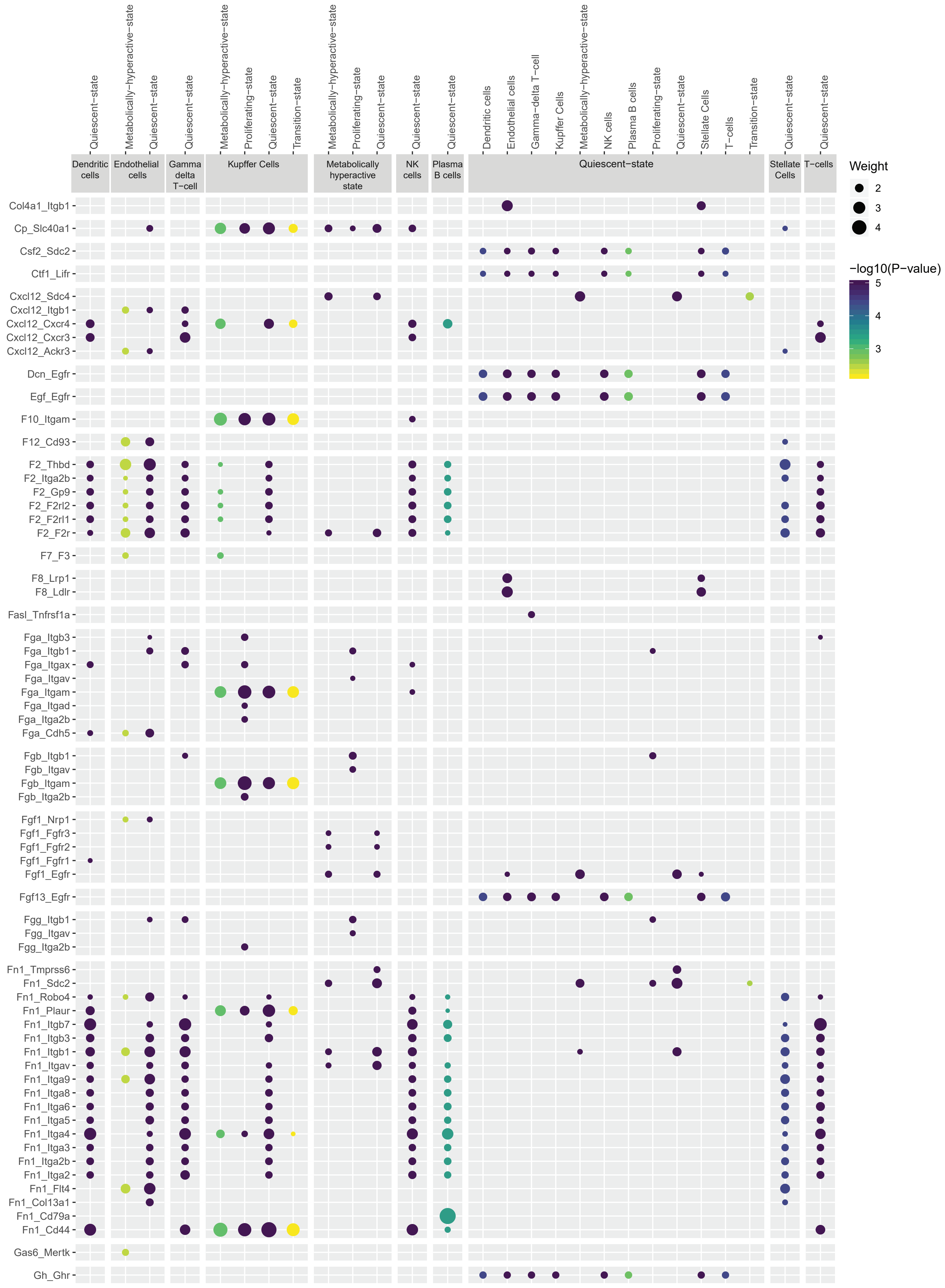

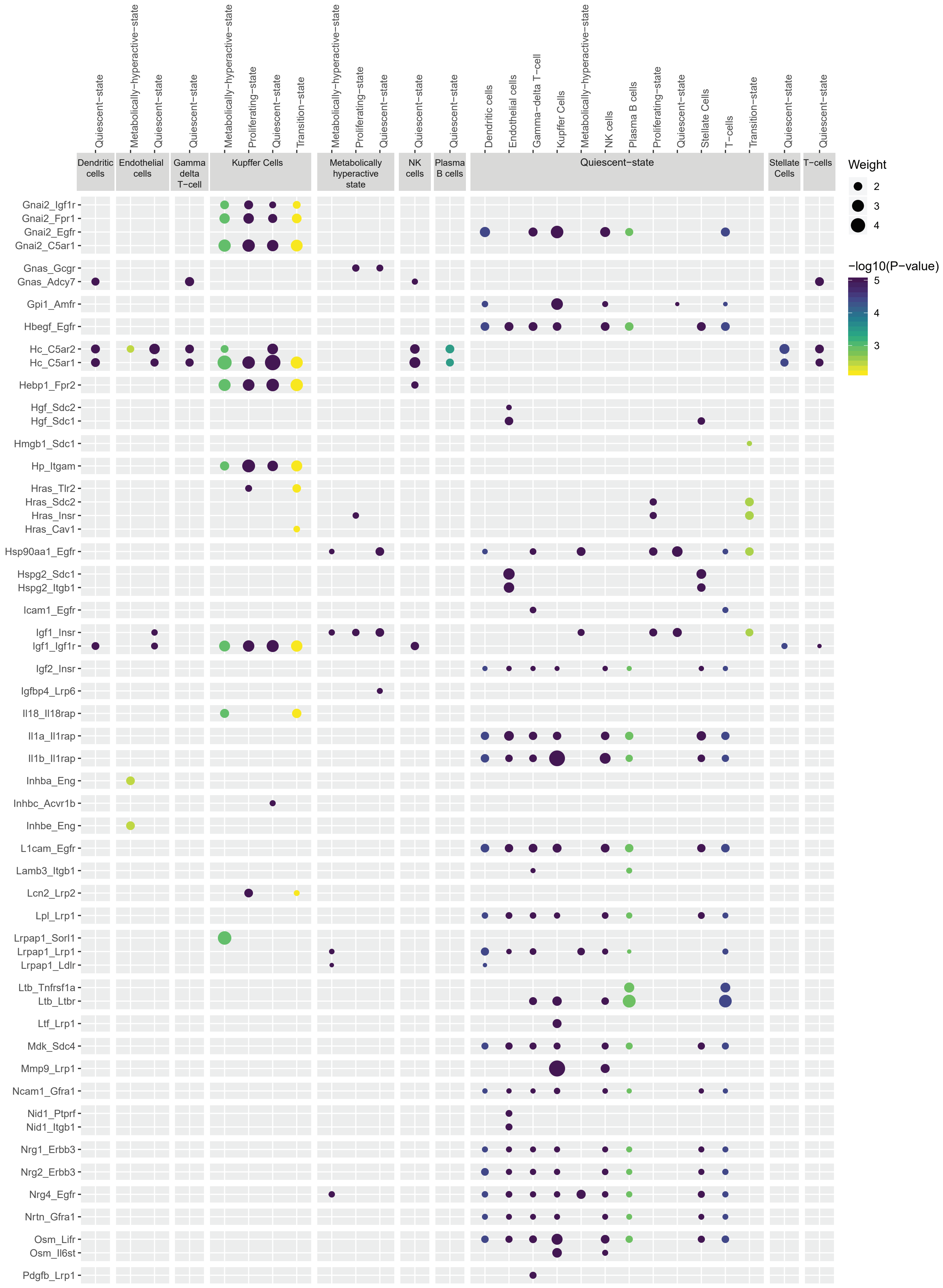

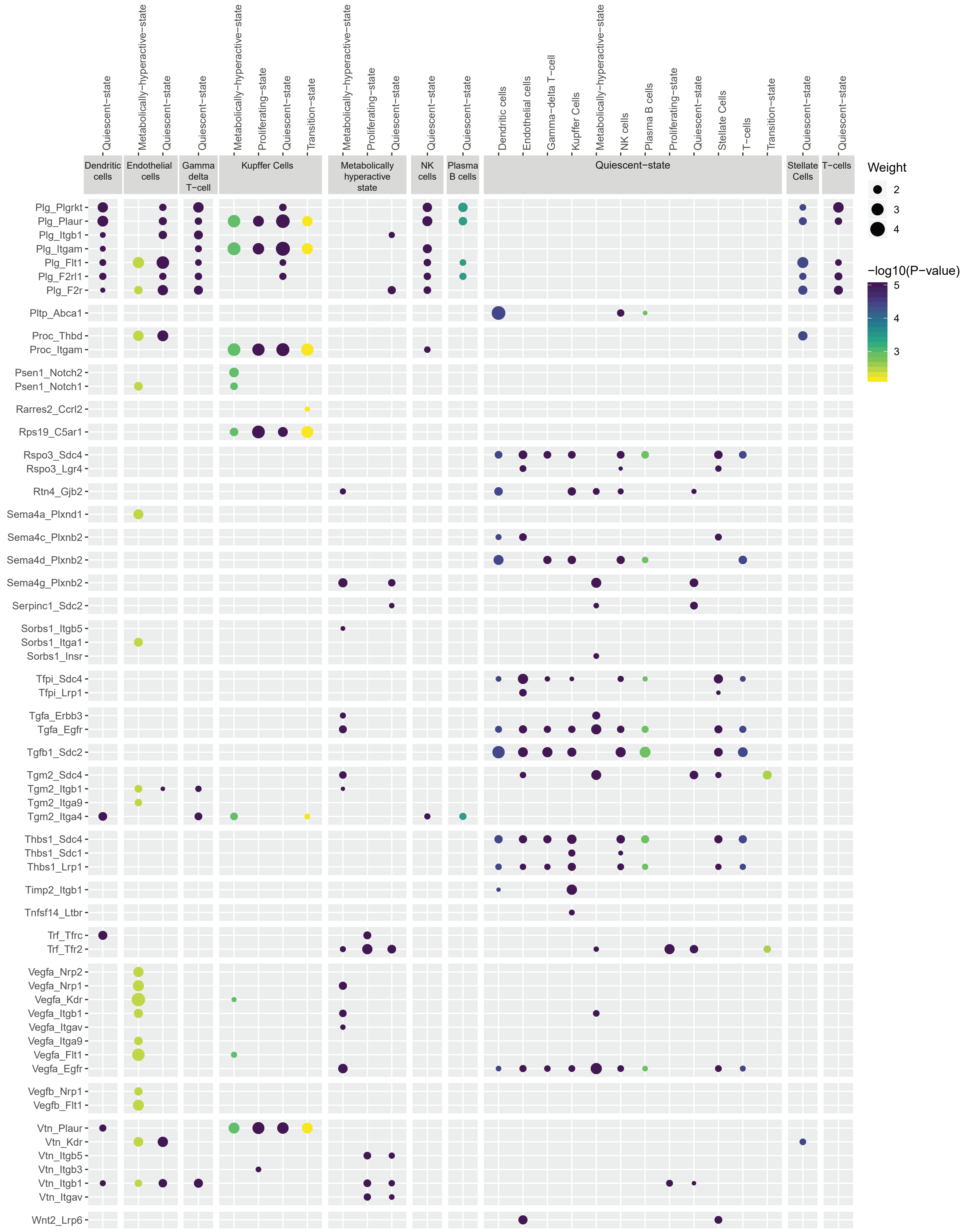

### Supplementary data file 4 PHx48_Ligand_receptor_interactions

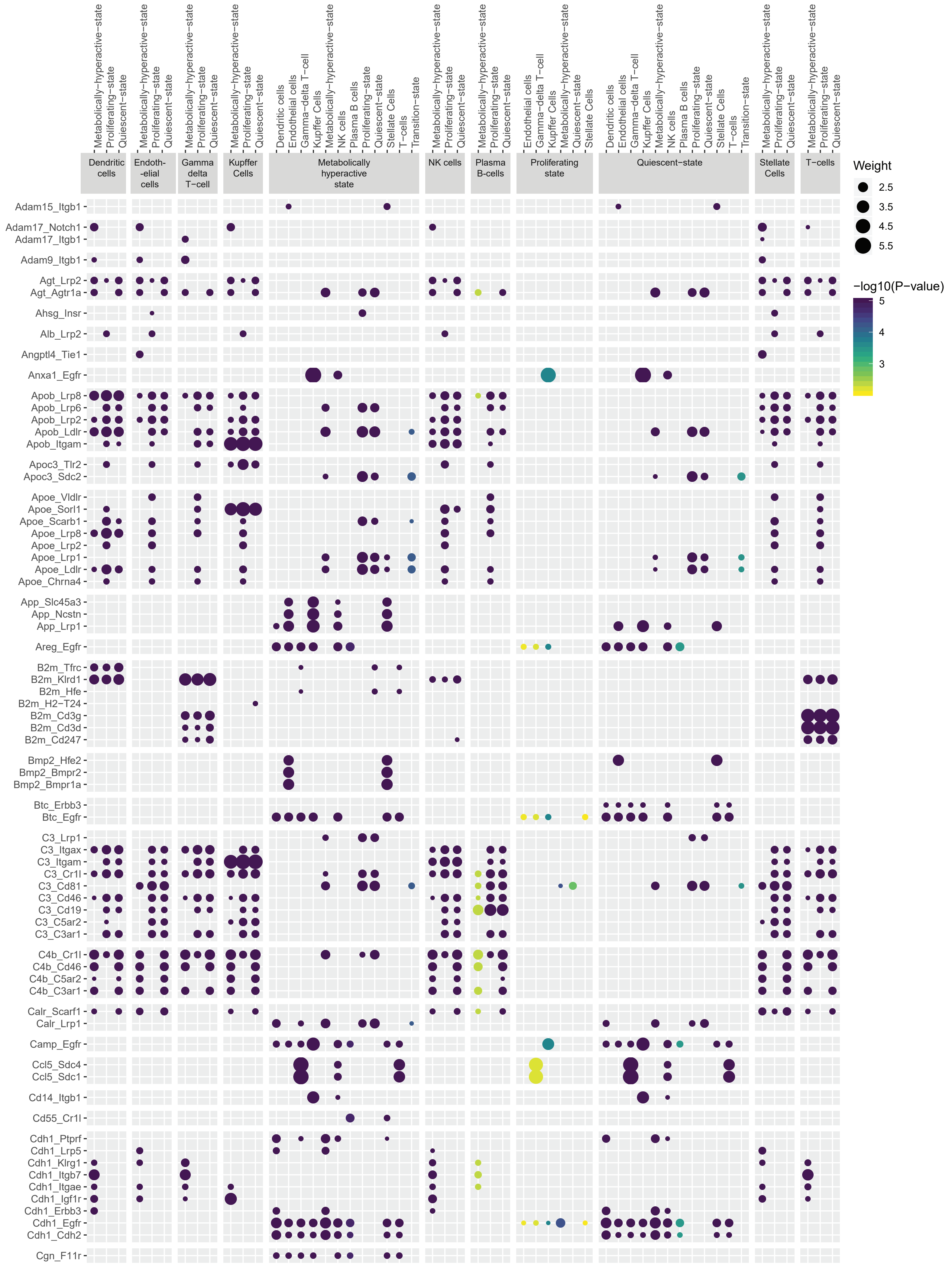

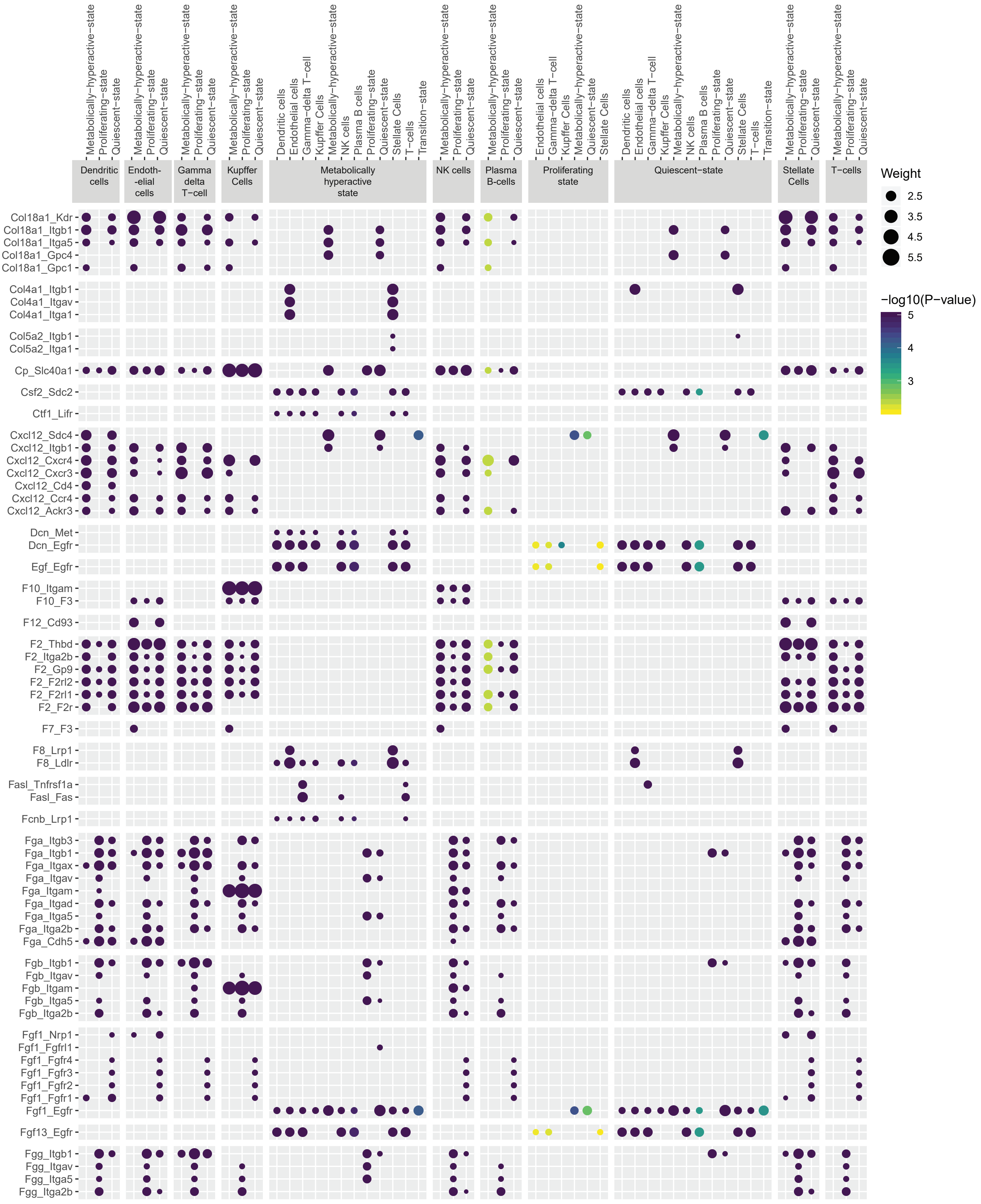

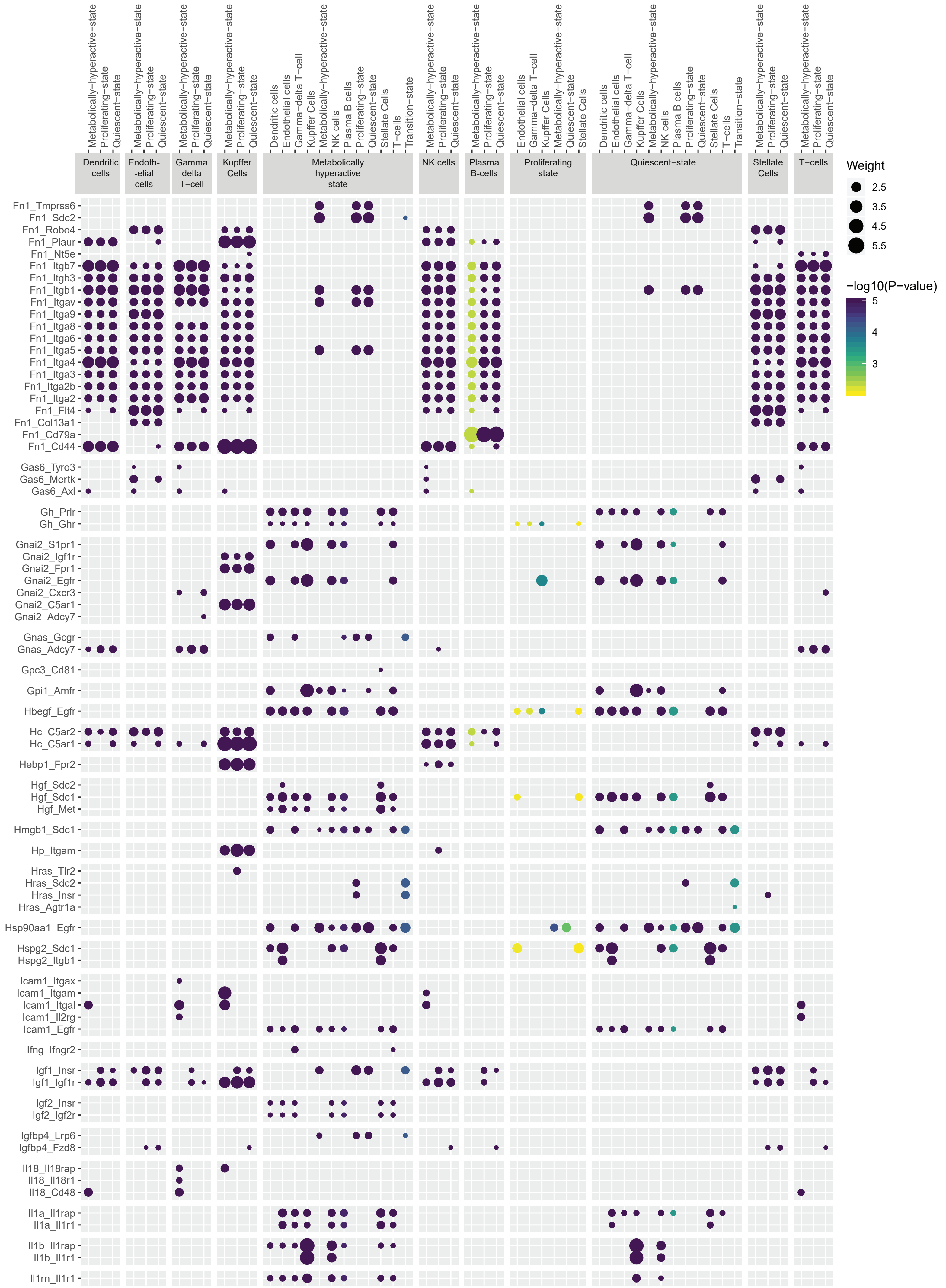

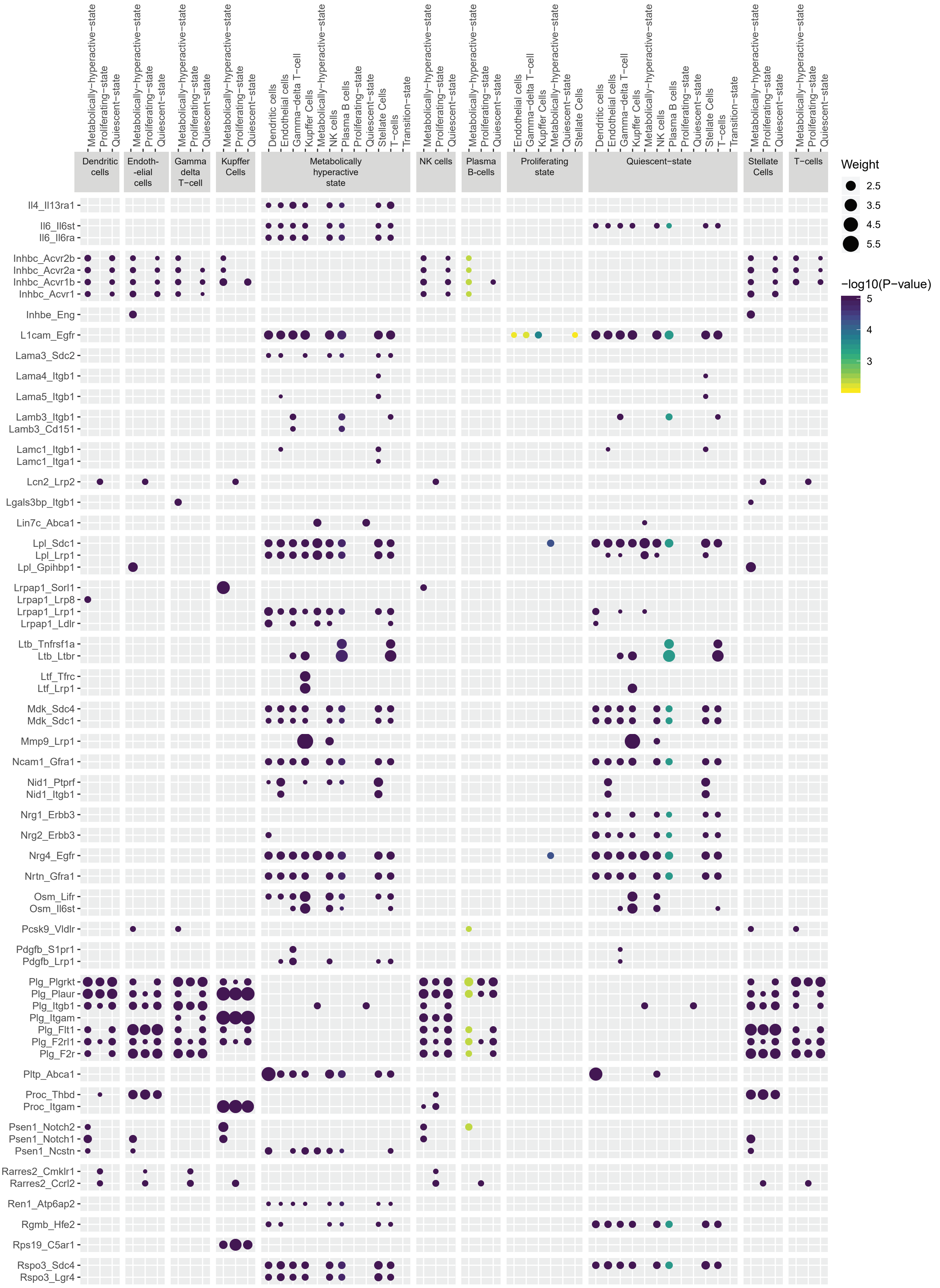

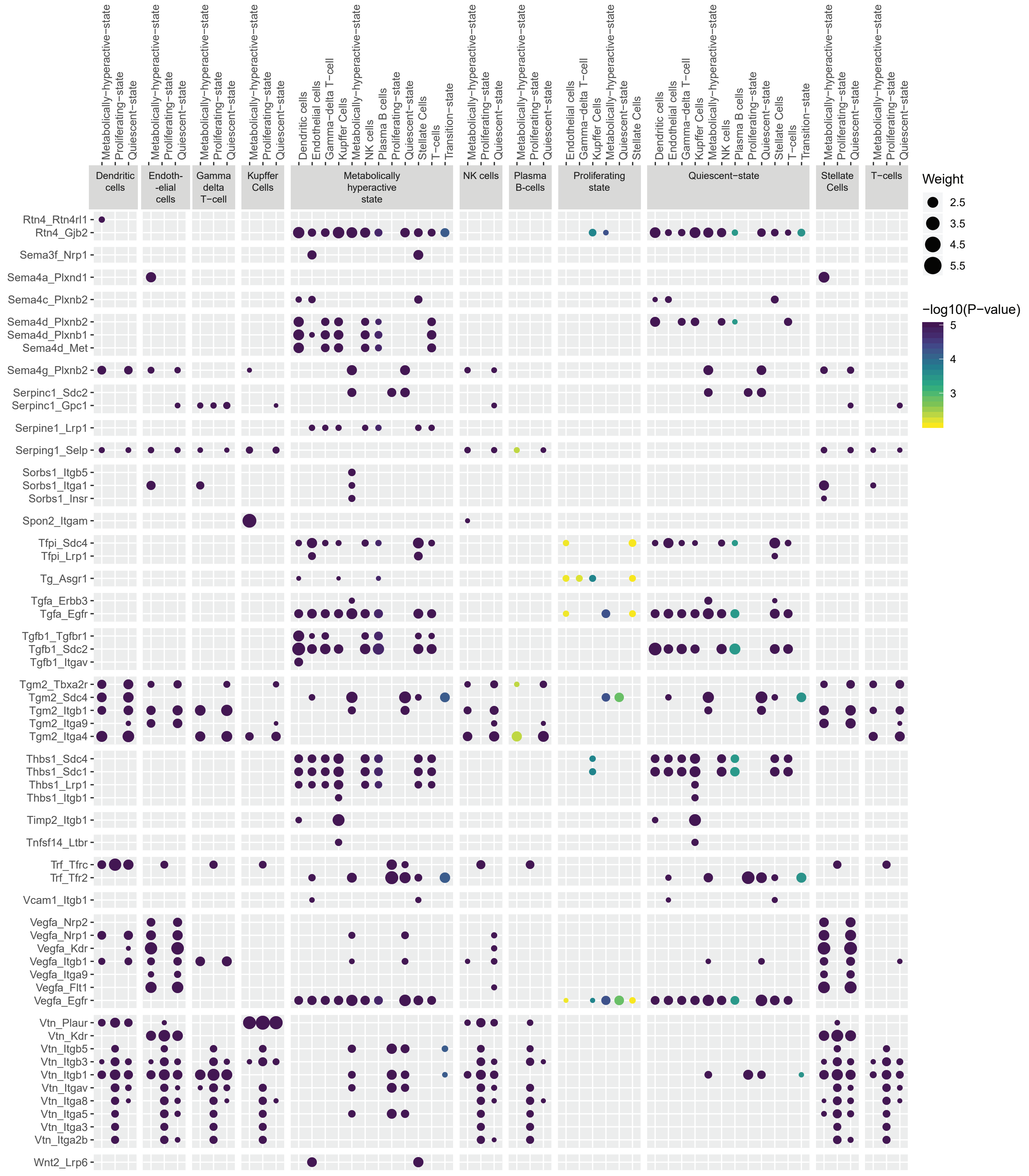

### Supplementary data file 5 PHx96__Ligand_receptor_interactions

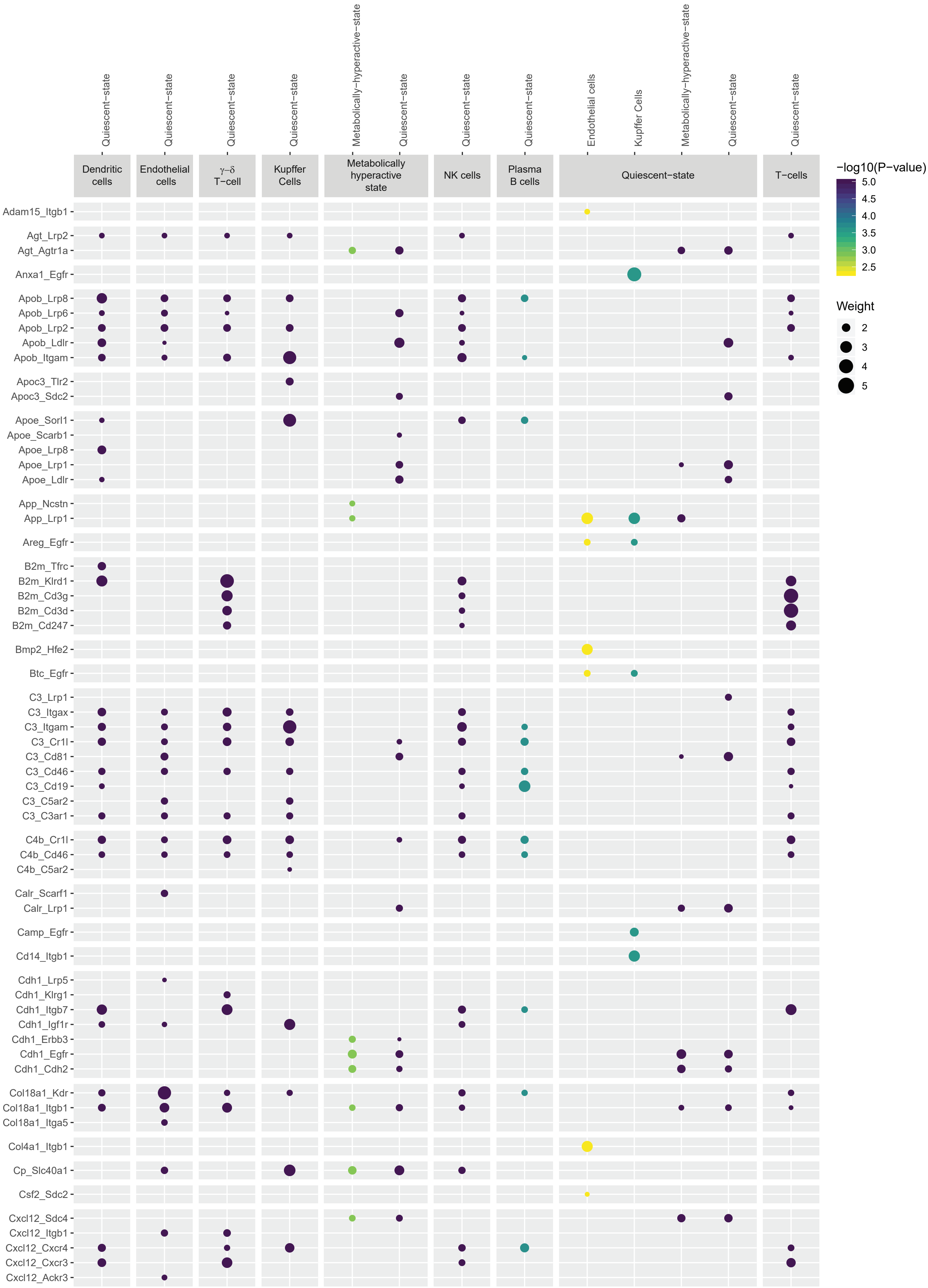

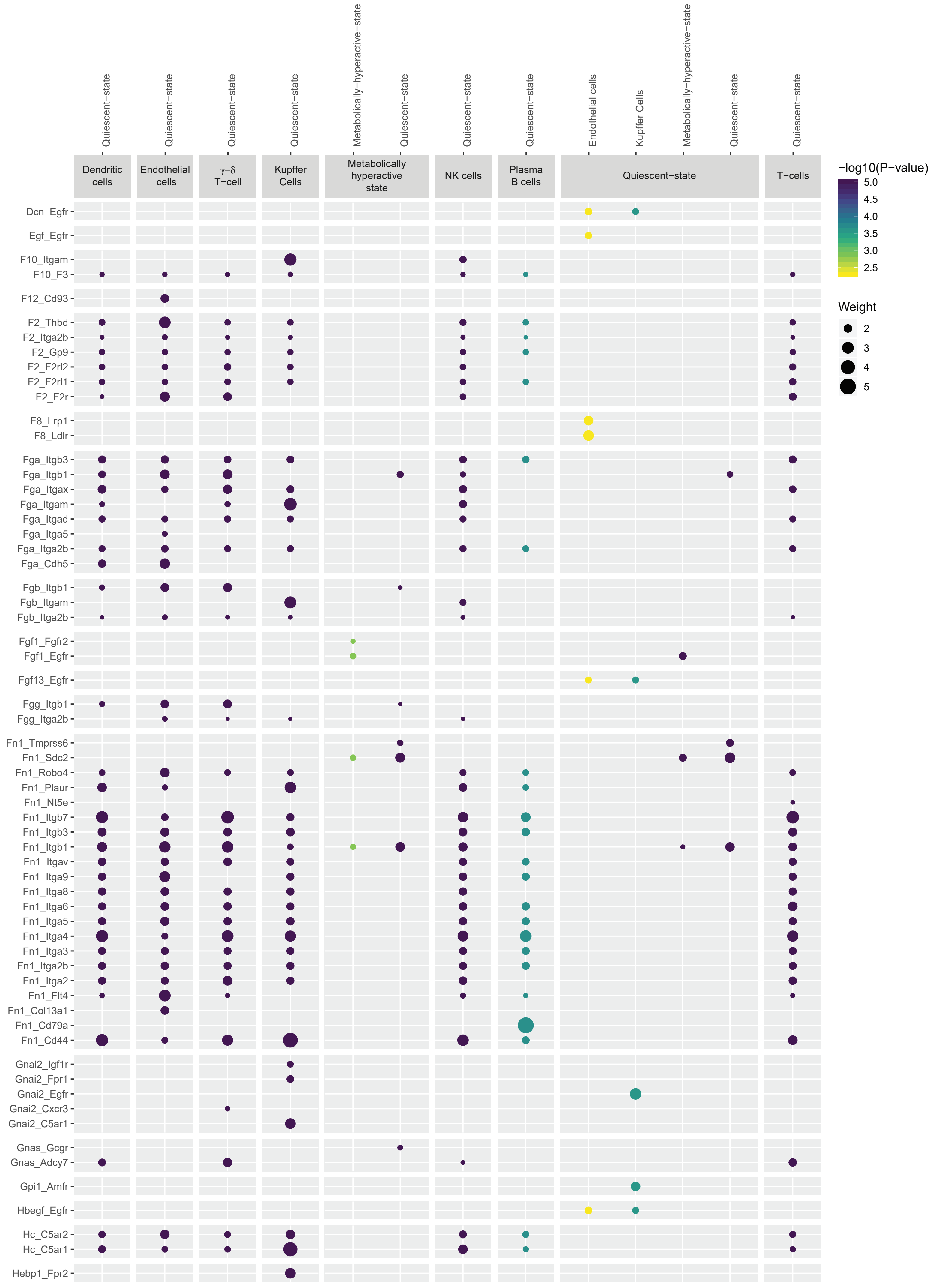

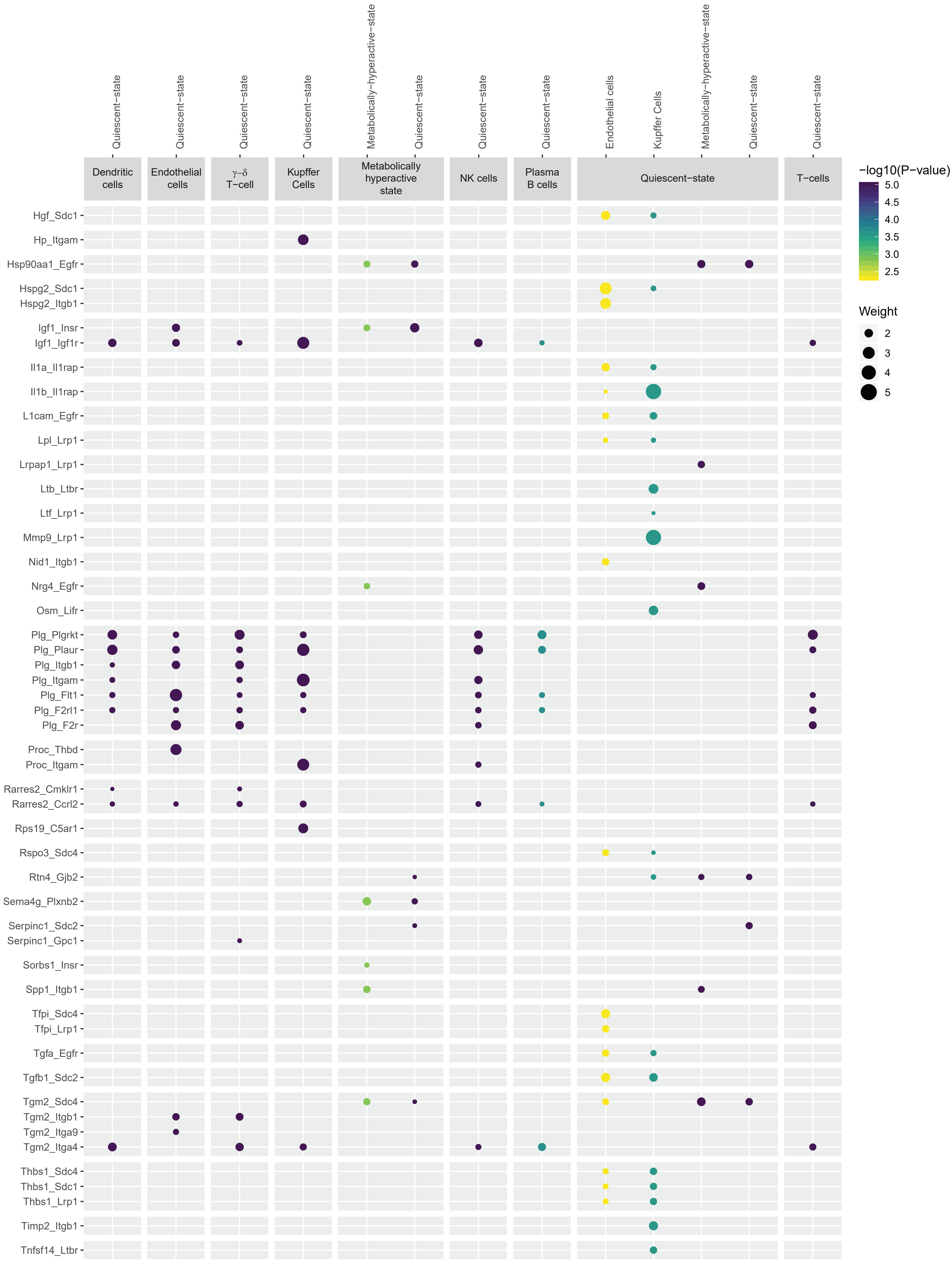

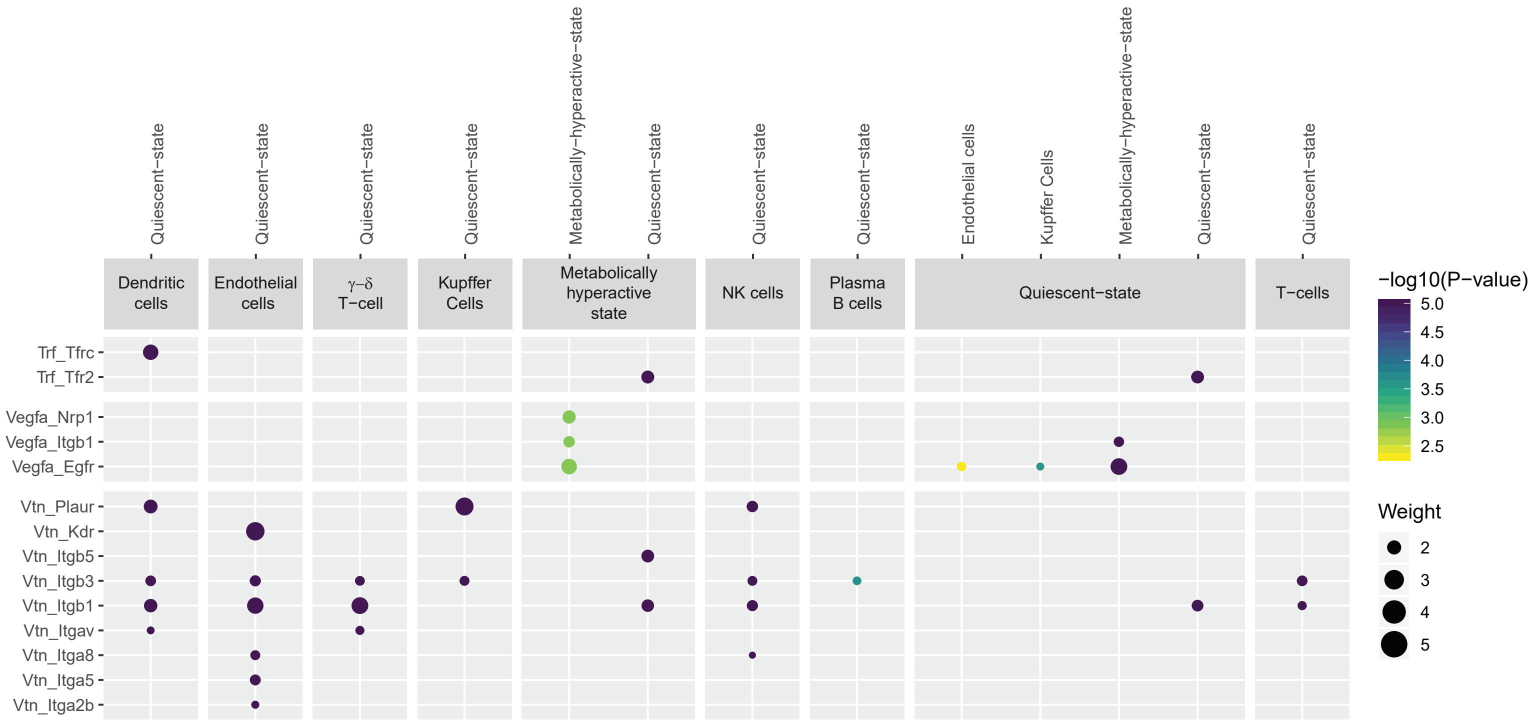
